# Orthogonal fluorescent chemogenetic reporters for multicolor imaging

**DOI:** 10.1101/2020.04.04.022111

**Authors:** Alison G. Tebo, Benjamien Moeyaert, Marion Thauvin, Irene Carlon-Andres, Dorothea Böken, Michel Volovitch, Sergi Padilla-Parra, Peter Dedecker, Sophie Vriz, Arnaud Gautier

## Abstract

Fluorescence microscopy is an indispensable tool in biological research, allowing sub-second and sub-micrometer mapping of molecules or processes inside living cells. Moreover, using spectrally separated fluorophores, one can observe multiple targets simultaneously, leading to a deeper understanding of the dynamic molecular interplays that regulate cell function and fate. Chemogenetic systems, which combine a protein tag and a synthetic fluorophore, provide certain advantages over fluorescent proteins since there is no requirement for chromophore maturation. However, the fluorophore promiscuity of chemogenetic systems renders two-color applications challenging. Here, we present the engineering of a set of spectrally orthogonal fluorogen activating tags based on the Fluorescence Activating and absorption Shifting Tag (FAST), that are compatible with two-color, live cell imaging. The resulting tags, greenFAST and redFAST, demonstrate orthogonality not only in their fluorogen recognition capabilities, but also in their one- and two-photon absorption profiles. A two-color cell cycle sensor based on greenFAST and redFAST is capable of detecting very short, early cell cycles in zebrafish development which had previously been difficult to image. Furthermore, this pair of orthogonal tags can be developed into split complementation systems that are capable of detecting multiple protein-protein interactions by live cell fluorescence microscopy.

## INTRODUCTION

Fluorescence imaging techniques allow one to follow the localization and activities of labeled biomolecules despite the crowded intracellular environment. The fluorescent labels can be entirely synthetic (e.g. organic fluorophores, quantum dots), entirely genetically encoded (e.g. fluorescent proteins) or a hybrid combination. While synthetic labels can be very bright and photostable, they are often difficult to target to a given biomolecule with high specificity. Genetically encoded fluorescent labels such as the green fluorescent protein (GFP)^1^ have become indispensable tools for biologists as they enable the facile generation of genetic fusions with virtually any protein of interest, however they suffer from slow (minutes to hours), oxygen-dependent maturation.

As an alternative, hybrid or chemogenetic systems have been proposed as a way to combine the advantages of synthetic labels^2^ with the targeting specificity of genetically encoded tags. Chemogenetic systems moreover often provide a great deal of experimental versatility through the ability to adapt the color of the fluorophore to the experimental conditions, simply by choosing a different cell-permeable and live-cell compatible molecule.

Chemogenetic systems can be generally classified by the nature of the interaction between the protein and the fluorophore. Halo-,^3^ SNAP-,^4^ and CLIP-tags^5^ are self-labeling tags that recognize their cognate ligands and catalyze their covalent attachment.^5^ In contrast, fluorogen activating proteins (FAPs) interact non-covalently with their cognate fluorogens to generate a fluorescent complex.^6^ Contrary to many self-labeling tags, the fluorogenic nature of these systems means that a fluorophore is initially in a non-fluorescent state, but become fluorescent upon binding. Separate washing steps to remove unbound dye are not required, and therefore dynamic processes can be followed more easily. A single FAP typically binds a variety of structurally similar fluorogens, providing a straightforward avenue to introduce color diversity by creating fluorogen derivatives. Multiple consecutive labeling steps with different fluorogens can also confer other benefits, such as enabling discrimination of the FAP moieties even in the presence of other fluorophores that emit in the same spectral region.^7^ However, the promiscuous nature of FAP binding severely limits multi-color imaging.

Our lab has recently developed a novel fluorogen activating protein, FAST (Fluorescence-Activating and absorption Shifting Tag), which is a 14 kDa monomeric protein that interacts rapidly and reversibly with a series of 4-hydroxybenzylidene rhodanine derivatives.^8^ The fluorogens that interact with FAST all do so with *K*_D_s in the micromolar to sub-micromolar regime and fluoresce from 540 nm to 600 nm, depending on the derivative.^7–9^ We recently expanded this system to create splitFAST, a split fluorescent reporter that displays rapid and reversible complementation, and that can readily be used as an indicator for molecular interactions.^10^ SplitFAST is unique for its reversibility and the kinetics of its association and dissociation. Like its ancestor FAST and related proteins, this split system shares the property of fluorogen promiscuity allowing facile adaptation of the emission wavelength to the experimental context.

While the fluorogen promiscuity has clear advantages in terms of spectral flexibility, it also makes it very difficult to use FAST or splitFAST to label two or more distinct targets with different colors, essentially rendering conventional multicolor imaging based on only these systems impossible. In fact, multicolor labeling with non-covalent fluorogen activating proteins presents a specific instance of a broader ligand-recognition problem: engineered proteins often recognize multiple ligands indiscriminately due to similarities in ligand chemical structure and binding mode.^11–13^ Many natural proteins, in contrast, exhibit exquisite sensitivity to highly related molecules with important biological consequences, as is the case for hormone receptors^14^ or cyclic nucleotide binding proteins^15,16^. Unraveling the principles underlying the selectivity of ligand binding is crucial not only for our understanding of native signaling processes, but also for drug design^17^ and synthetic biology^18^.

Here, we present the development of orthogonal, color-selective FAST variants for multicolor imaging. The resulting variants show orthogonality in their selectivity for a particular fluorogen, resulting in labels that show selective green or orange/red emission. We demonstrate the usefulness of our constructs for both one and two-photon excitation fluorescence microscopy, two-color super-resolution imaging, and fluorescence lifetime imaging. We also demonstrate two-color fluorescence microscopy in both eukaryotic cell culture and zebrafish models, which allow the observation of very short cell cycles early in zebrafish development. Finally, we generated reversible split fluorescent reporters for the simultaneous detection of two transient protein-protein interactions. Our work supports that competitive selection schemes in directed evolution can help provide insights into selectivity as well as accelerate the development of novel systems.

## RESULTS AND DISCUSSION

We chose to focus our efforts on the combination of FAST with the fluorogen HMBR, which forms a green fluorescent complex with a *K*_D_ of 0.1 μM, and HBR-3,5DOM, which forms an orange-red fluorescent complex with a *K*_D_ of 1 μM. Both fluorogens are structurally similar, differing only by their substituents modifying the 4-hydroxybenzyl moiety (3-methyl vs. 3,5-dimethoxy) (**Figure 1a**). As in many fluorogenic hybrid systems,^19,20^ these small substitutions allow the generation of various colors and thus many hybrid systems are comprised of a suite of structurally related fluorogens that interact with a single, genetically-encoded tag.

**Figure 1.**
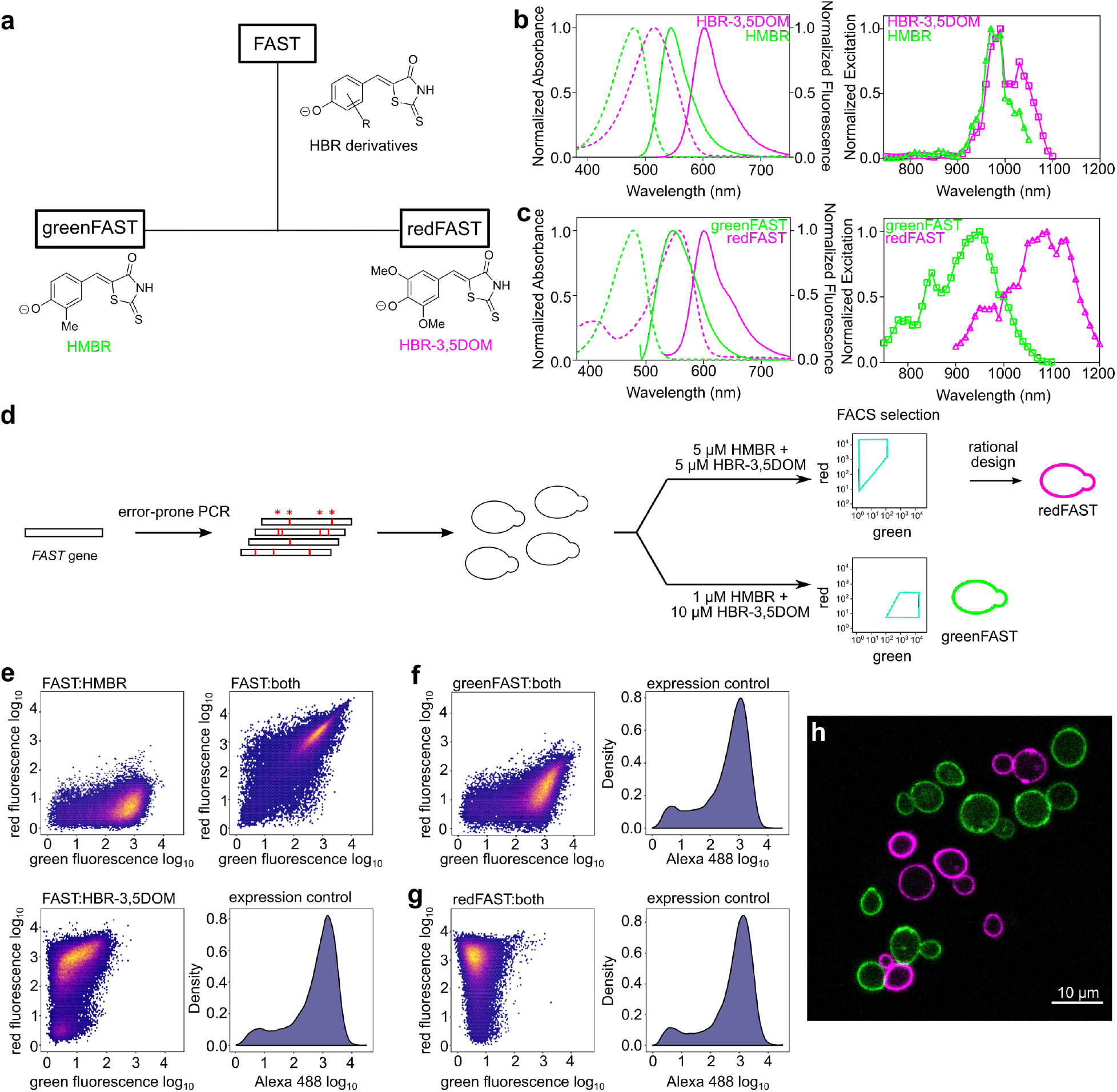
Engineering and properties of greenFAST and redFAST. **(a)** FAST promiscuously binds HBR derivatives while greenFAST and redFAST were evolved to bind selectively HMBR or HBR-3,5DOM. **(b) (left)** Absorbance (dotted lines) and emission (solid lines) spectra of iFAST:HMBR (green) and iFAST:HBR-3,5DOM (magenta). **(right)** Two-photon excitation spectra of iFAST:HMBR (green) and iFAST:HBR-3,5DOM (magenta). **(c) (left)** Absorbance (dotted lines) and emission (solid lines) spectra of greenFAST:HMBR (green) and redFAST:HBR-3,5DOM (magenta). **(right)** Two-photon excitation spectra of greenFAST:HMBR (green) and redFAST:HBR-3,5DOM (magenta). **(d)** Selection and design strategy for spectrally orthogonal FAST systems. **(e-g)** Yeast cells expressing FAST (**e**), redFAST (**f**) and greenFAST (**g**) were analyzed by flow cytometry in the presence of either only HMBR (5 μM) or HBR-3,5DOM (10 μM) or in the presence of both (5 μM HMBR and 10 μM HBR-3,5DOM). Efficient induction of protein expression was verified through independent labeling with an Alexa488-conjugated antibody**. (h)** Confocal micrographs of a mixture of yeast cells expressing greenFAST (green) and redFAST (magenta) in the presence of 5 μM HMBR and 10 μM HBR-3,5DOM. Scale bar 10 μM.

We set out to develop orthogonal FAST:fluorogen systems with selectivity for HMBR or HBR-3,5DOM using a directed evolution strategy. A random library of FAST variants (10^6^ mutants) was developed using error-prone PCR; yeast surface display coupled with FACS sorting allowed us to select for variants with orthogonal fluorogen selectivity (**Figure 1d**). To select for HMBR-selective ‘green’ variants, FACS was performed in the presence of 1 μM HMBR and 10 μM HBR-3,5DOM, conditions in which most yeast cells are doubly labeled, given the *K*_D_s of 0.1 μM for HMBR and 1 μM for HBR-3,5DOM observed with FAST. The ‘greenest’ cells in these conditions were selected to enrich for clones that preferentially bind HMBR. The selection of HBR-3,5DOM-selective ‘red’ variants was more challenging given the higher affinity of FAST for HMBR, so FACS was performed in the presence of 5 μM HMBR and 5 μM HBR-3,5DOM. These conditions allowed us to select for cells expressing clones with higher preference for HBR-3,5DOM despite competition with HMBR. Five rounds of selection were performed for the green (selective for HMBR) and red (selective for HBR-3,5DOM) orthogonal variants, respectively.

After the fifth round of selection, twenty-four clones were picked randomly and screened for selectivity using flow cytometry and sequenced. Nine individual sequences were found in the selection for the green variants, with certain sequences representing a high proportion of the total sequences (7 times)(**Table S1**). The same analysis was performed for the red variants, resulting in eight individual sequences with certain ones also being highly represented (7 or 6 times) (**Table S2**). Five of the nine green clones and all eight red clones were subcloned into a bacterial expression vector for purification and screening *in vitro*. The *in vitro* screening revealed that the green variants retained their affinity for HMBR while losing their affinity for HBR-3,5DOM, in accordance with a gain of selectivity for HMBR (**Table S1**). Most dramatically, the green clones 1 and 6 displayed an over 10-fold lower affinity for HBR-3,5DOM compared to native FAST (**Figure S1a**). Conversely, the red variants all retained their ability to bind HBR-3,5DOM while losing an order of magnitude in affinity for HMBR (**Figure S1a**, **Table S2**).

The selectivity of a given construct can be estimated using the ratio of the binding constants of each fluorogen (*K*_D,HBR-3,5DOM_/*K*_D,HMBR_ for the green variants and *K*_D,HMBR_/*K*_D, HBR-3,5DOM_ for the red ones), which should be larger than ten for ensuring selectivity. The clones isolated by FACS screening for HBR-3,5DOM were not selective, despite displaying a 10-fold reduction in HMBR binding affinity. Indeed, the resulting equivalence of the two binding affinities (around 1 μM) resulted in a ratio of binding constants of ~1, and was thus insufficiently selective.

To screen for improved selectivity, highly repeated mutations isolated from the different clones were combined to try to introduce potential additive effects. Four new variants were generated for the green system, while three were developed for the red system (**Figure S1b**, **Table S3**). However, none of the new green variants displayed brighter complexes with HMBR. Clone 1 from the original selection was retained and renamed greenFAST, which displays *K*_D_s of 0.09 μM and 16.2 μM for HMBR and HBR-3,5DOM, respectively (**Table 1**, **Figure S2**). In the case of greenFAST, the three mutations present (G21E, P68T, G77R) confer selective binding of HMBR over HBR-3,5DOM. We similarly introduced two additional mutations (F28L and E46Q) that needed to be added to the red clone 10 so as to confer selectivity for HBR-3,5DOM, in order to generate a highly selective variant that displays a 100-fold lower affinity for HMBR as compared to FAST and a 10-fold higher selectivity for HBR-3,5DOM vs HMBR (*K*_D_s of 12 μM and 1.2 μM for HMBR and HBR-3,5DOM, respectively) (**Table 1**, **Figure S2**). This variant, now possessing a total of five mutations (F28L, E46Q, R52A, E81V, S99N), was retained and renamed redFAST. The mutation F28L reduced the affinity of the protein for both fluorogens (**Table S3**), possibly through repacking of the N-terminal domain against the β sheet, which has previously been shown to modify the p*K*_A_ of the chromophore of PYP.^21^ However, subsequent addition of E46Q rescued the affinity for HBR-3,5DOM, leaving the reduced affinity for HMBR relatively untouched and resulting in a favorable ratio *K*_D,HMBR_/*K*_D,HBR-3,5DOM_ of 10 (**Table S3**).

**Table 1.**
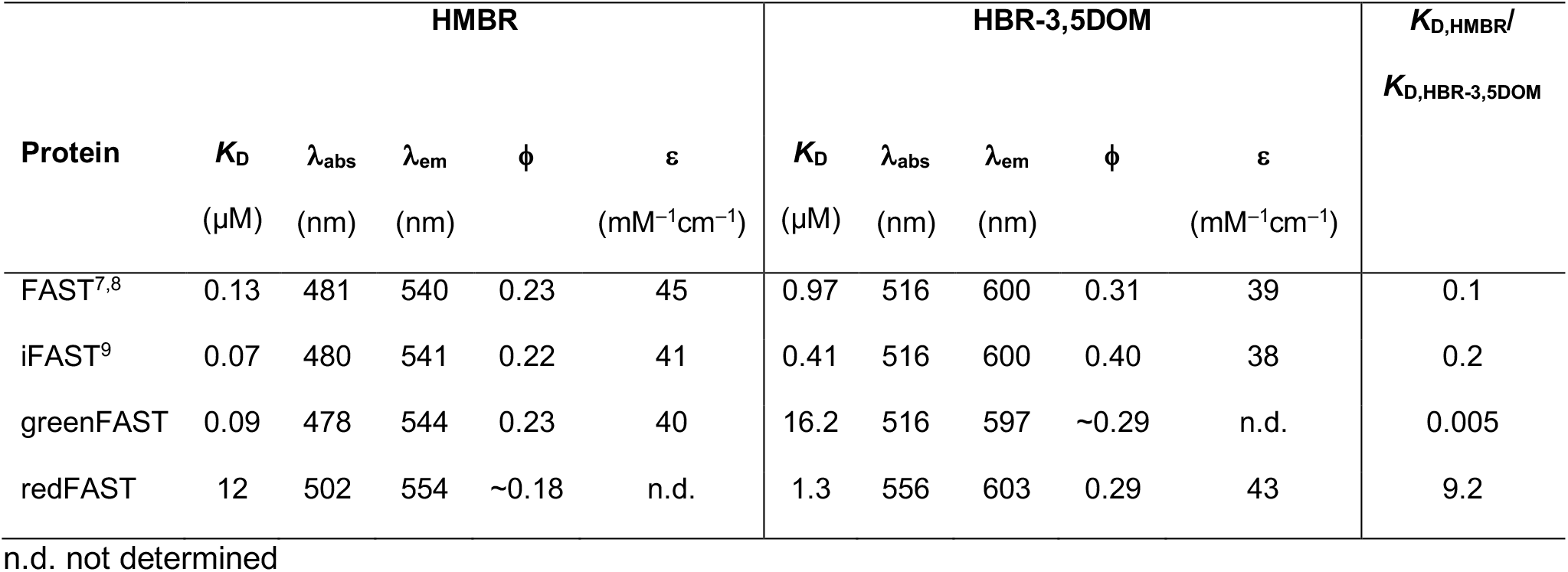
Physicochemical characteristics of FAST, iFAST, greenFAST, and redFAST with HMBR and HBR-3,5DOM in PBS pH 7.4. Abbreviations are as follows: λ_abs_, wavelength of maximal absorption; λ_em_, wavelength of maximal emission; *ε*, molar absorptivity at λ_abs_; ϕ, fluorescence quantum yield; *K*_D_ thermodynamic dissociation constant.

To assess the orthogonality of the system, yeast cells expressing FAST, greenFAST, and redFAST at the cell surface were analyzed by flow cytometry (**Figure 1e-g**). FAST-expressing clones in the presence of 5 μM HMBR and 10 μM HBR-3,5DOM predictably bind both fluorogens and display both green and red fluorescence (**Figure 1e**). In comparison, under the same conditions greenFAST and redFAST show remarkably similar fluorescence profiles to FAST labeled with only one of these fluorogens, demonstrating their selective binding profiles (**Figure 1f,g**). Two-color imaging using confocal microscopy allowed us to easily distinguish the two cell populations, visually confirming the flow cytometry results (**Figure 1h**). The successful engineering of these orthogonally selective tags demonstrates the strength of a competitive selection scheme, which enabled us to identify key mutations governing the affinity of the protein scaffold for a particular fluorogen. Given the structural similarity of the two ligands, a wholly rational approach to design ligand specificity would be impeded by the difficulty of identification of distal interactions that affect binding *a priori*.

A more detailed examination of the absorption and emission spectra of greenFAST and redFAST revealed an unexpected spectral orthogonality, which improves their performance for two-color applications. HBR-3,5DOM complexes with FAST and iFAST (an improved FAST variant) are most efficiently excited at 516 nm and exhibit broad absorption spectra^7,9^ (**Figure 1b**), making cross-excitation by a 488 nm laser possible. Thus, a system that is simply orthogonal in terms of binding affinity could still suffer from crosstalk excitation of both fluorogens at the same wavelength. However, the absorption of redFAST is redshifted by nearly 40 nm, placing it squarely in a spectral region where excitation by 543 nm and 561 nm lasers is more optimal (**Figure 1c**). This shift can be ascribed to the mutations of E46 and R52, two residues that are known for their role in hydrogen bonding to the chromophore in PYP, the parent protein of FAST. Mutational analysis in PYP has shown that these mutations result in a red-shifted PYP chromophore absorption spectrum similar to what we observed in redFAST.^21^ It is worth noting that at least one of these two mutations is present in every clone isolated after FACS screening, which is likely due to the selective pressure imposed by the laser and filter sets used for screening. Indeed, while the absolute molecular brightness of redFAST is lower than that of both iFAST:HBR-3,5DOM and mCherry, the good spectral match in excitation wavelength means that redFAST is 1.7 brighter than iFAST^9^ and 1.2 times brighter than mCherry^22^ when excited at 561 nm. Interestingly, this trend extends to two-photon excitation. The two photon excitation spectra of iFAST with HMBR and HBR-3,5DOM display strong overlap (**Figure 1b**), while the spectra of redFAST and greenFAST are red-shifted and blue-shifted, respectively, resulting in orthogonal excitation profiles in two-photon mode (**Figure 1c**).

The selectivity and spectral orthogonality of greenFAST and redFAST make them ideal for two color microscopy applications. To test their performance as cellular markers, greenFAST and redFAST were fused to a number of proteins with various subcellular localizations (**Figure 2a-e**). Co-expression of greenFAST and redFAST fusions localized to different subcellular structures in mammalian cells demonstrated that the two proteins could be used together for two-color imaging applications and that fusion of the tags did not interfere with the expected localization. Photobleaching measurements revealed that redFAST exhibited greater photostability than FAST or iFAST, while greenFAST was, on the contrary, more sensitive to photobleaching (**Figure S3**, **SI Text 1**).

**Figure 2.**
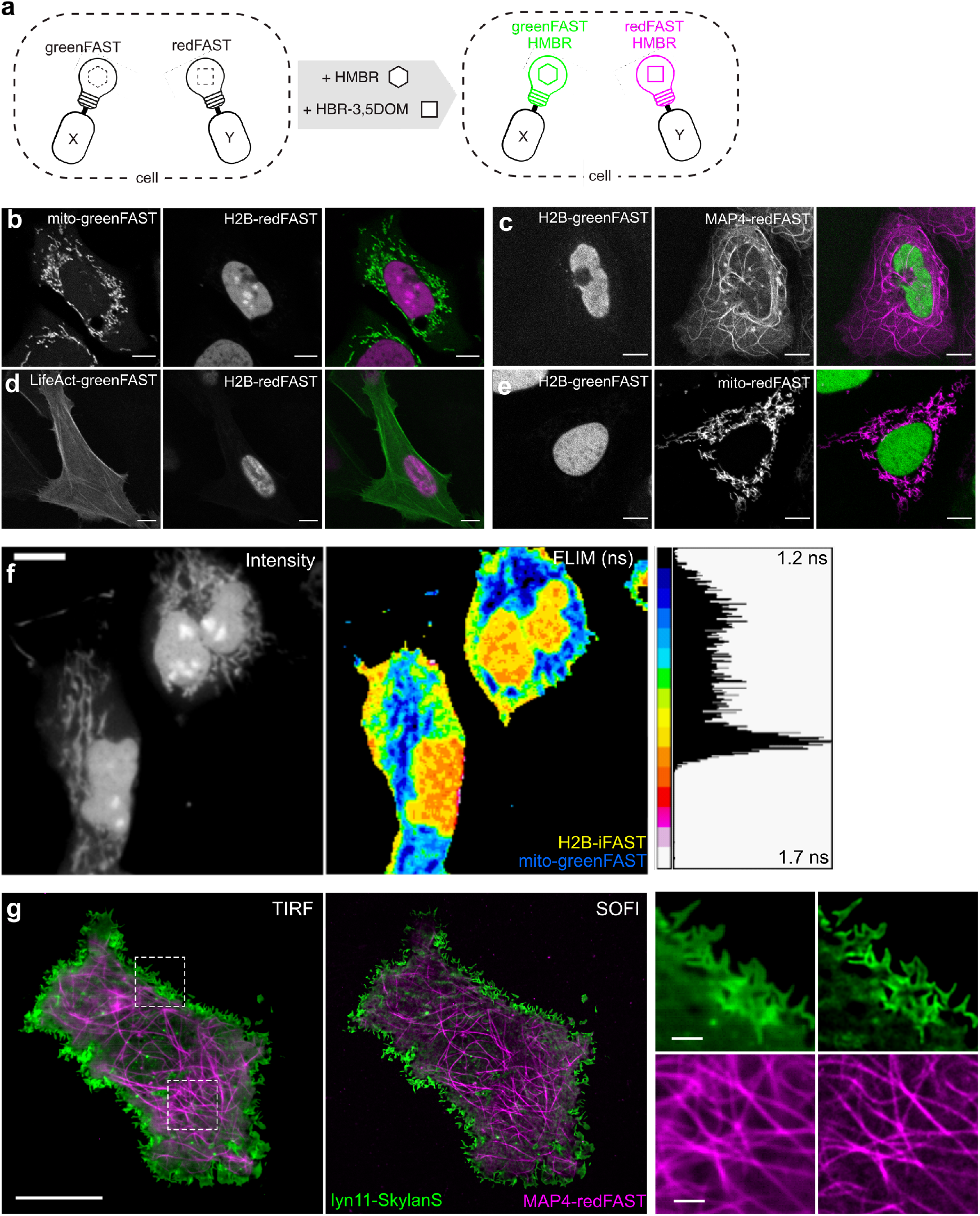
Imaging of greenFAST and redFAST in live cells. **(a-e)** Two-color imaging of greenFAST and redFAST fusions in cells. **(b)** Micrograph of U2OS cells expressing mito-greenFAST and H2B-redFAST. **(c)** Micrograph of U2OS cells expressing H2B-greenFAST and MAP4-redFAST. **(d)** Micrograph of U2OS cells expressing LifeAct-greenFAST and H2B-redFAST. **(e)** Micrograph of U2OS cells expressing H2B-greenFAST and mito-redFAST. **(b-e)** Cells were labeled with 5 μM HMBR and 10 μM HBR-3,5DOM. Scale bars 10 μM. **(f)** COS-7 cells expressing mito-greenFAST and H2B-iFAST labeled with 5 μM HMBR. Scale bars 5 μM. **(g)** Averaged TIRF (left) and pcSOFI (right) images of COS-7 cells expressing lyn11-Skylan-S and MAP4-redFAST. Cells were labeled with 5 μM HBR-3,5DOM. Scales bars 10 μM (left) and 1 μM (right).

We also characterized the excited-state lifetimes of redFAST, greenFAST, and iFAST using fluorescence lifetime imaging microscopy (FLIM)^23^ on live COS-7 cells expressing H2B fusion proteins. Similar to many fluorescent proteins, the fluorescence decays of FAST-based complexes are best fit with a biexponential function (**Tables S4-S10**). The slow lifetime component of redFAST:HBR-3,5DOM was similar to that of iFAST:HBR-3,5DOM (2.48 ± 0.07 ns n = 17 vs. 2.77 ± 0.05 ns, n = 15) (**Table S4**). In contrast, the greenFAST:HMBR and iFAST:HMBR complexes displayed distinguishable slow lifetime components (1.18 ± 0.09 ns, n = 15 vs. 1.70 ± 0.02 ns, n = 20), (**Table S4**), which could be used to selectively image both labels in live cells using FLIM (**Figure 2f**). Furthermore, imaging greenFAST and redFAST in the presence of both fluorogens did not change the measured lifetime.

Finally, we tested the performance of greenFAST and redFAST for one modality of super-resolution fluorescence microscopy, namely super-resolution optical fluctuation imaging SOFI^24–26^. SOFI provides diffraction-unlimited spatial resolution by relying on the analysis of spontaneous single-molecule ‘blinking’ of the fluorophores, and can operate under a broad range of conditions using any single-molecule sensitive wide-field microscope. COS-7 cells expressing greenFAST and stained with 5 μM HMBR did not show appreciable single-molecule intensity fluctuations and thus did not yield insightful images. However, redFAST did show single-molecule intensity fluctuations suitable for SOFI imaging. Two-color SOFI imaging could be achieved using live mammalian cells co-expressing microtubule-targeted redFAST (stained with HBR-3,5DOM) and membrane-targeted Skylan-S,^27^ a green fluorescent protein that was showed to be robustly well-performing in pcSOFI^28^ (**Figure 2g**). Second-order pcSOFI analysis showed the background rejection and two-fold gain in spatial information intrinsic to the method.^29^ This result illustrates redFAST’s fitness for multicolor SOFI.

Green/red FAST labelling seems particularly promising for dynamic recording *in vivo*. We thus decided to apply greenFAST and redFAST to the FUCCI (fluorescence ubiquitination cell cycle indicator) technology.^30^ FUCCI is a technique for delineating cell cycle phases via the use of a red and a green fluorescent protein fused to the N-terminal domains of Cdt1 and geminin, two cell cycle regulators whose levels show biphasic cycling during the cell cycle. However, the analysis of very short cell cycles as found in early fish or amphibian embryos is not possible due to the time limitations imposed by the slow maturation of typical GFP-like fluorescent proteins. For example, the zFUCCI system adapted to zebrafish, though better at delineating the G1/S transition, was not able to track cell cycle phases earlier than the 6 somite stage (12 hpf).^31^ A more recent improvement of the FUCCI system^32^ allowed analysis of the shortest cell cycles in mouse embryonic stem cells dividing in culture, lasting 9-10 h. These advances still precluded the analysis of the very fast cell cycles occurring in zebrafish early embryos, where cells may divide every 15-18 min^33–35^ before the mid-blastula transition (MBT) and for which G1 and G2 pattern and tempo of appearance are still unknown.

A stable mammalian cell line expressing a FAST-based FUCCI, consisting of redFAST fused to the N-terminal domain of Cdt1 and greenFAST fused to the N-terminal domain of geminin, enabled the observation of multiple cell cycles over long time-lapse acquisitions (24-28 hrs) (**Figure 3a-d**). We thus decided to use the same ubiquitin ligase domains which had previously proved useful in zebrafish, zCdt1(1-190) and zGeminin(1-100) and fused them with greenFAST and redFAST, respectively. The mRNA coding both proteins was injected into a zebrafish embryo at the one-cell stage and time-lapse imaging was performed starting from 256-cell stage embryos with one image every 5 minutes. The first cycle gave faint signals, as expected for very short cycles with quite reduced G1 or G2 phases, but the two following cycles (9^th^ and 10^th^) gave excellent signals allowing individual nuclei to be monitored over time (**Figure 3e,f**, **Movie S1**). The 9^th^ cell cycle displayed a clear G1/S transition and lasted 15 min. The 10^th^ cycle was longer (30 min) and a well-demarcated G2 phase appeared. Indeed, all phase transitions (M/G1, G1/S, S/G2 and G2/M) were clearly visible in our system, which benefits from the absence of fluorescence maturation in greenFAST and redFAST because of the almost-instantaneous fluorogen binding. Interestingly, zCdt1(1-190) is targeted for degradation by CUL4,^31^ which is normally turned-on and turned-off at the beginning and end of S phase.^32^ This should give zCdt1(1-190) a biphasic accumulation regime (first during G1 phase, with an abrupt disappearance during S phase, then during G2), which is indeed what was observed during the 10^th^ cycle (**Figure 3e,f**). It is also noteworthy that during this cycle, the G1/S, S/G2 and G2/M transitions were all perfectly synchronous in the two sister cells (**Figure 3e,f**).

**Figure 3.**
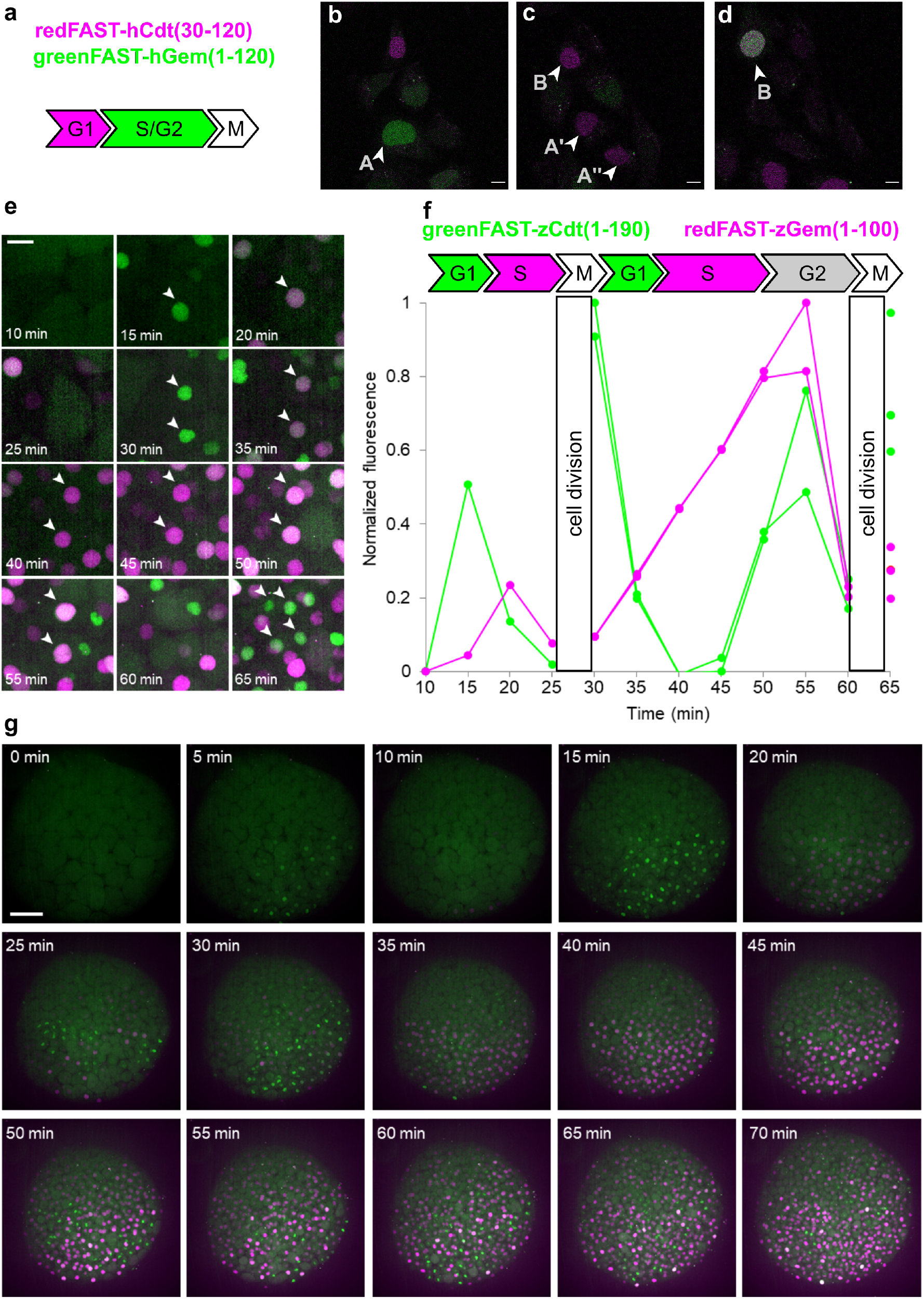
Cell cycle sensors based on orthogonal FASTs. **(a-d)** U2OS cells stably expressing a FUCCI cell cycle sensor. **(a)** Design of a mammalian cell cycle sensor. Tracking of individual cell cycles is possible through stable expression of redFAST-hCdt(30-120) and greenFAST-hGem(1-120). **(b-d)** Cell A can be tracked S/G2 (**b**, 40 mins) through division (**c**, 6 hrs 10 mins) while Cell B (**c**, top arrow) can be tracked through the G1-S transition (**d**, 19 hrs 50 mins). Images were taken every 5 mins (see also **Movie S1**). Cells were maintained at 37°C and 5% CO_2_ and were imaged in HEPES-buffered DMEM without phenol red and supplemented with 5 μM HMBR and 10 μM HBR-3,5DOM. Scale bars 10 μM. **(e-g)** Zebrafish embryos were injected with redFAST-zGem(1-100)-P2A-greenFAST-zCdt1(1-190) mRNA at one-cell stage, and time-lapse imaging was performed starting from 256-cell stage on embryos incubated with 5 μM HMBR and 5 μM HBR-3,5DOM. **(e)** Single cell visualization at higher magnification. Scale bar 20 μM. **(f)** Corresponding quantification of fluorescence signal over time. **(g)** Whole embryo imaging (see also **Movie S2**). Scale bar 100 μm.

In addition, fluorescence recording over the whole embryo revealed proliferation patterns and asynchronic division as early as 256 cells (**Figure 3g**, **Movie S2**). These patterns were previously described in either fixed embryos^36^ or in live embryos analyzed by label-free non-linear microscopy, though these did not give access to individual phases of the cell cycle^35^. Our approach is the first dynamic study of all phase transitions in the cell cycles at the mid-blastula transition in zebrafish embryogenesis,^37^ highlighting the strength of multicolor chemogenetic reporters with rapid labeling kinetics. Indeed, reaching the detailed analysis of cell cycles as short as 15 min (as compared to 9-10 h in recent studies^32^) and as early as 3.5 hpf (as compared to 12 hpf as reported earlier^31^) is only possible by using fluorescent labels with quasi-instantaneous fluorescence maturation as the ones developed here.

SplitFAST is the only reversible fluorescence complementation reporter with rapid association and dissociation kinetics.^10^ We reasoned that greenFAST and redFAST could also be used for the design of split reporters, provided that they retained these characteristics as well as their orthogonality, which would open the possibility for the readout of multiple interactions. Especially promising in this regard would be the facile readout of multiple or sequential protein interactions as the appearance of fluorescence combined with the reversibility inherent to the splitFAST system provide good contrast and temporal resolution, the combination of which is currently impossible to achieve.

GreenFAST and redFAST were split into N-terminal and C-terminal fragments at the same site used to create splitFAST from FAST and fused to the FK506-binding protein (FKBP) and the FKBP-rapamycin-binding domain of mammalian target of rapamycin (FRB) that interact together in the presence of rapamycin. All the mutations that confer selectivity to greenFAST and redFAST occur in the N-terminal fragments (named greenNFAST and redNFAST respectively), meaning that the C-terminal fragment (named CFAST11) is the same for the two split systems. Split-greenFAST and split-redFAST were assessed directly in mammalian cells by confocal microscopy. We measured the association of the split constructs by inducing the formation of heterodimers of FRB and FKBP with rapamycin (**Figure 4a,b,e**). Upon addition of 500 nM rapamycin, split-redFAST and split-greenFAST rapidly reassembled into their cognate fluorogen:protein complex in the presence of both fluorogens with an average 6- and 8-fold increase in fluorescence (**Figure 4e**). Furthermore, the fluorescence time course revealed that split-greenFAST and split-redFAST show similarly rapid kinetics (seconds-minutes) as has previously been observed for splitFAST (**Figure 4a,b**).

**Figure 4.**
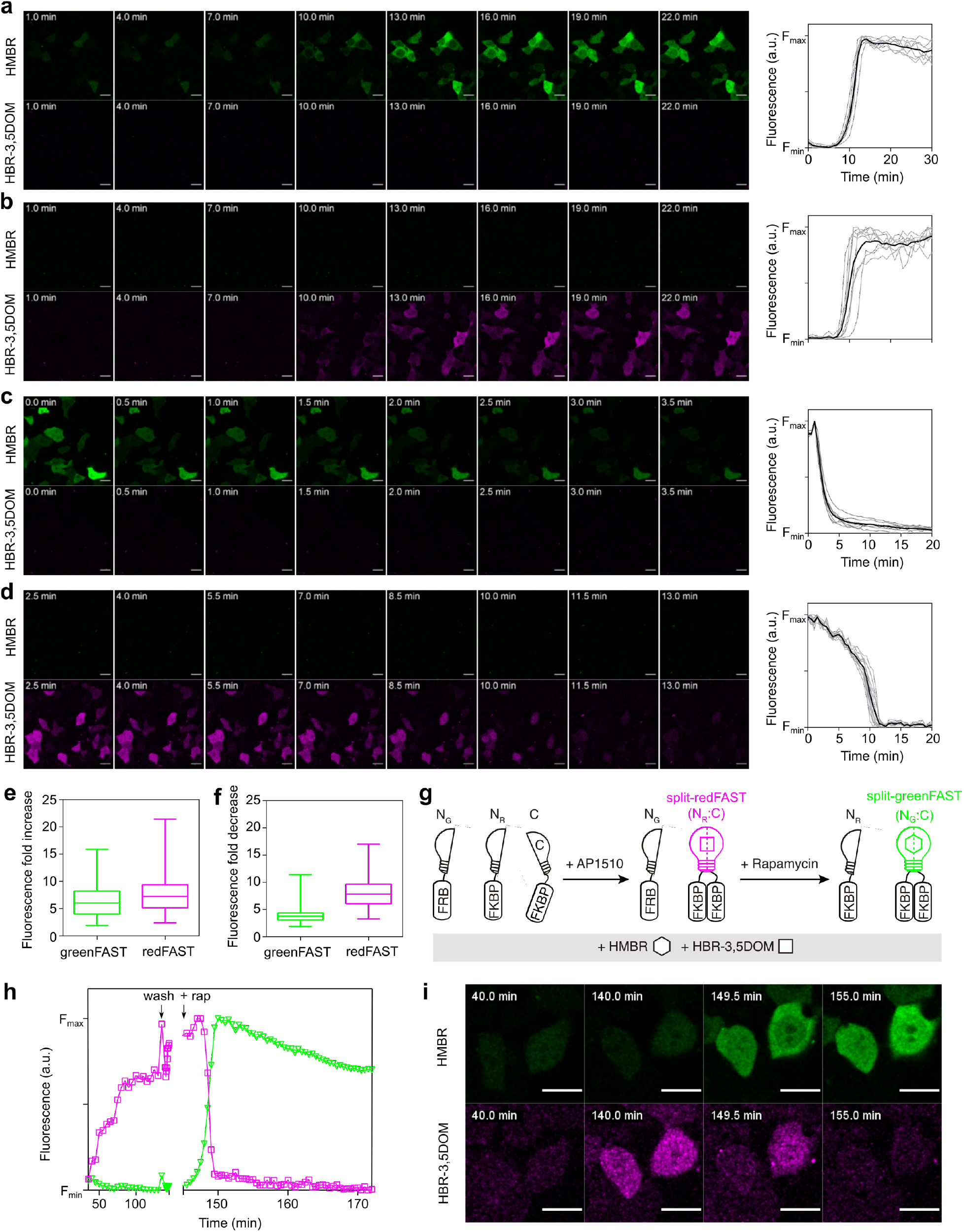
Orthogonal reporters of protein-protein interactions. **(a,b)** HEK293T cells co-expressing FK506-binding protein (FKBP) fused to CFAST11 and FKBP-rapamycin-binding domain of mammalian target of rapamycin (FRB) fused to either greenNFAST **(a)** or redNFAST **(b)** were labeled with both 5 μM HMBR and 10 μM HBR-3,5DOM, and imaged before and after addition of 500 nM rapamycin. The green channel shows HMBR fluorescence, while the magenta channel shows HBR-3,5DOM fluorescence. Graphs show the temporal evolution of the fluorescence intensity after rapamycin addition. **(c,d)** HEK293T cells co-expressing FKBP fused to CFAST11 and FBBP fused to either greenNFAST **(c)** or redNFAST **(d)** treated with 100 nM AP1510 and labeled with both 5 μM HMBR and 10 μM HBR-3,5DOM. Cells were then imaged before and after the addition of 1.1 μM rapamycin. The green channel shows HMBR fluorescence, while the magenta channel shows HBR-3,5DOM fluorescence. Graphs show the temporal evolution of the fluorescence intensity after rapamycin addition. **(e)** Fluorescence fold increase upon FRB-FKBP association for split-greenFAST and split-redFAST. Box and whiskers (max, min). Data from three independent experiments. Number of cells, green = 157, red = 125. **(f)** Fluorescence fold decrease upon FKBP-FKBP dissociation for split-greenFAST and split-redFAST. Box and whiskers (max, min). Data from three independent experiments. Number of cells green = 174, red = 135. **(g-i)** Detection of two protein-protein interactions. HEK293T cells co-expressing FKBP-CFAST1, FKBP-redNFAST and FRB-greenNFAST were labeled with both 5 μM HMBR and 10 μM HBR-3,5DOM, and image before and after addition of 500 nM rapamycin. The formation of FKBP-FKBP homodimers detected by redFAST was induced by addition of AP1510. Addition of rapamycin leads to the concomitant dissociation of FKBP-FKBP homodimer and formation of FKBP-FRB. **(h,i)** Representative traces and images showing the evolution of the fluorescence signals of split-redFAST and split-greenFAST during the experiment. Scale bars 10 μm.

In order to assess the reversibility of the complex, N-greenFAST (resp. N-redFAST) and CFAST11 were fused to FKBP. The disruption of an AP1510-mediated FKBP-FKBP homodimer by rapamycin was followed as a decrease in fluorescence (**Figure 4c,d,f**). The dissociation of split-greenFAST and split-redFAST upon rapamycin addition was likewise characterized by rapid kinetics similar to those observed for splitFAST(**Figure 4f**).

To test whether these two split systems can be used to measure two protein-protein interactions in the same cell, we monitored the exchange from FKBP-FKBP homodimer to FRB-FKBP heterodimer. Split-redFAST was used to detect the FKBP-FKBP homodimer, which was then dissociated by addition of rapamycin, triggering the concomitant association of FKBP and FRB, detected with split-greenFAST (**Figure 4g,h,i**). The association and dissociation of the two complexes occurred in the same kinetic regimes (seconds-minutes) as what has previously been observed with splitFAST for each interaction separately (**Figure 4h,i**, **Movie S3**). Both greenFAST and redFAST share a common C-terminal fragment when split, which enables the detection of switch-like protein-protein interactions that share a common partner, but also does not preclude the detection of two separate interactions with two C-terminal fragments fused to separate proteins.

In conclusion, we have developed two closely related chemogenetic systems that allow facile multicolor genetically-encoded labelling of living systems. Our systems offer a set of unique advantages that complement the suite of current imaging techniques, such as the straightforward observation of transient or rapidly cycling processes enabled by the absence of delays linked to chromophore maturation. They also display favorable optical properties for both one- and two-photon imaging and more advanced modalities such as FLIM or super-resolution imaging. We also showed that these proteins could be used for biosensing, by developing split reporters that report *in situ* interactions via rapid and reversible complementation, while offering straightforward readout by monitoring the appearance of fluorescence emission. We expect that our systems will strongly expand the range of questions that can be answered using fluorescence imaging.

## SUPPORTING INFORMATION

Supporting information contains Supplementary Movies S1-S3, Supplementary Figures S1-S3, SI Text 1, Supplementary Tables S1-S12, and the Materials and Methods section.

## NOTES

The authors declare the following competing financial interest: A.G. is co-founder and hold equity in Twinkle Bioscience / The Twinkle Factory, a company commercializing the FAST technology.

## ACKNOWLEDGEMENT

We thank K. D. Wittrup, for providing us with the pCTCON2 vector and the EBY100 yeast strain for the yeast display selection. We also thank the flow cytometry facility CISA (Cytométrie Imagerie Saint-Antoine) of UMS LUMIC at the Faculty of Medicine of Sorbonne University, and, more particularly, Annie Munier for her assistance. This work has been supported by the European Research Council (ERC-2016-CoG-724705 FLUOSWITCH to A.G., ERC StG 714688 NanoCellActivity to P.D., and ERC-2019-CoG-863869 FUSION to S.P-P.), the Wellcome Trust Core Award (203141 to S.P-P), and the Research Foundation-Flanders (G0B8817N to P.D. and 1514319N to B.M.)

