## Supplementary information file for "Orthogonal fluorescent chemogenetic reporters for multicolor imaging"

**Contents**

Supplementary Figures S1-3

Supplementation Movie Legends S1-3

SI Text 1

Supplementary Tables S1-12

Materials and Methods

Supplementary References

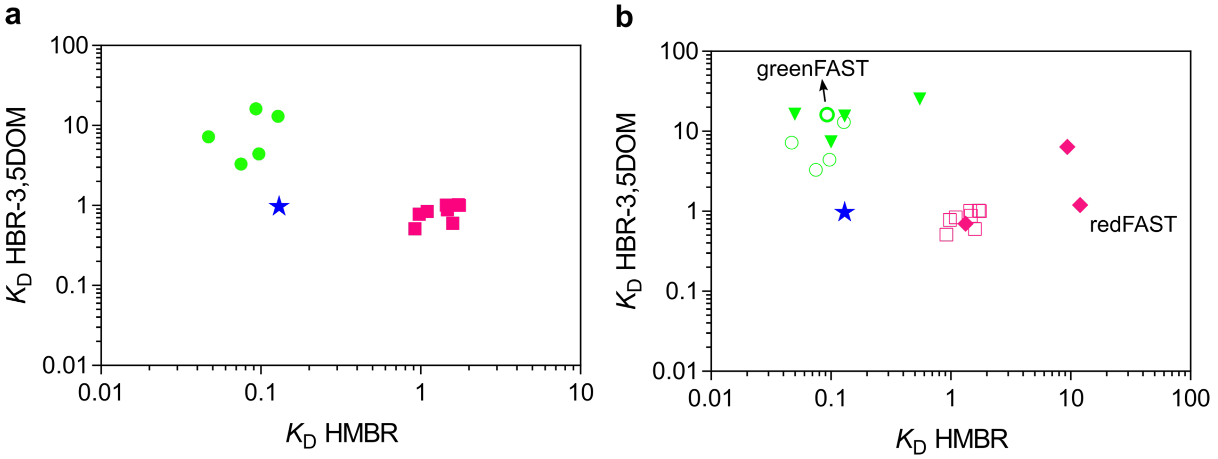


**Figure S1.** Comparison of the $K_D$s for HMBR and HBR-3,5DOM of FAST (blue star) and clones selected from FACS (a) and variants constructed through rational design (b, filled markers). Green dots: greenFAST selection, magenta dots: redFAST selection.

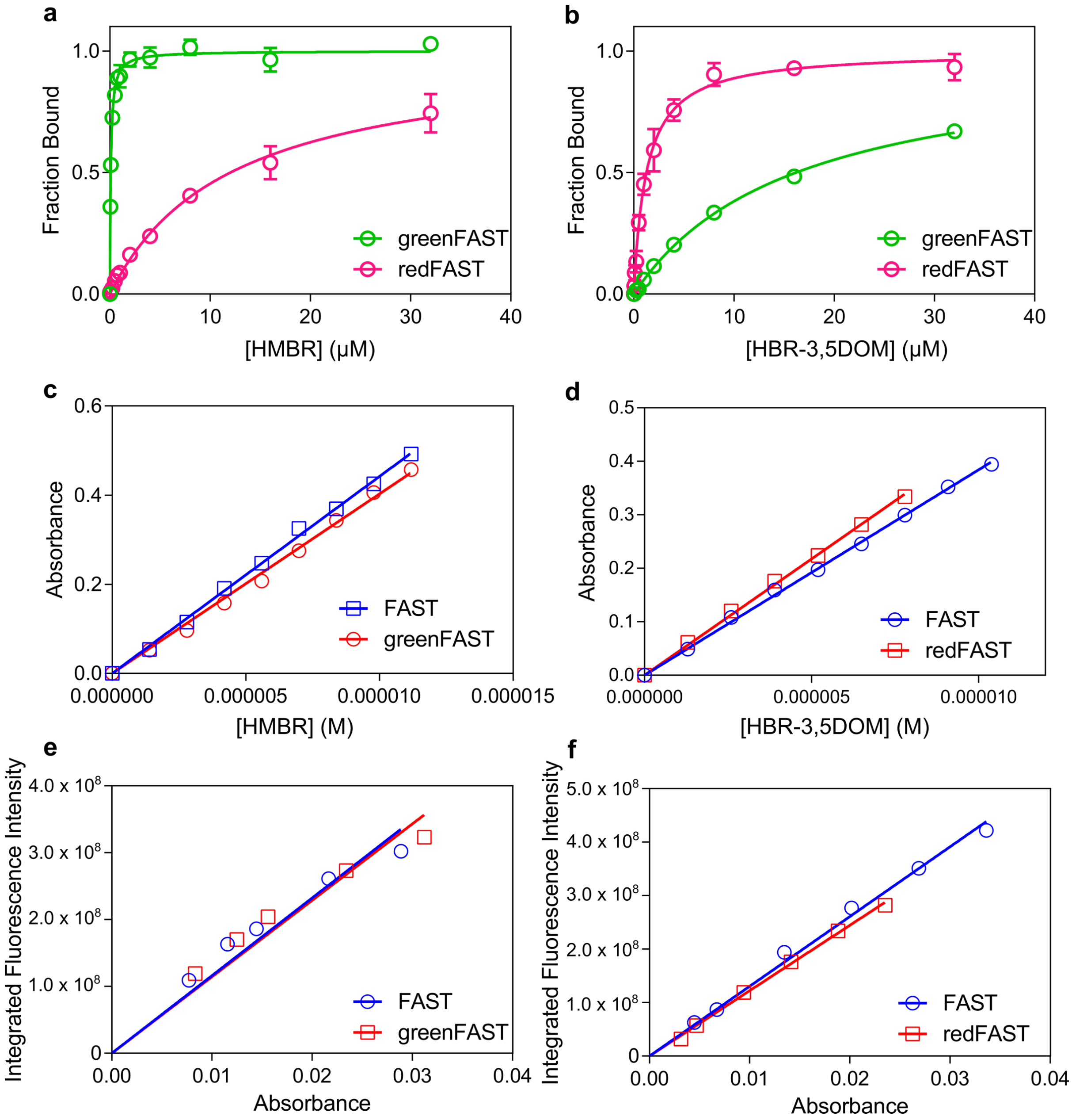


**Figure S2. (a,b)** Affinities of greenFAST and redFAST for a) HMBR and b) HBR-3,5DOM. Mean of n = 3, represented as mean ± sem, protein concentration 100 nM. **(c,d)** Determination of molar absorptivity for c) greenFAST and d) redFAST with their cognate fluorogen by forward titration and standardization with FAST (protein concentration, 40 µM). **(e,f)** Determination of quantum yield for e) greenFAST and d) redFAST with their cognate fluorogen by reciprocal dilution using FAST:fluorogen as a standard (protein concentration, 40 µM).

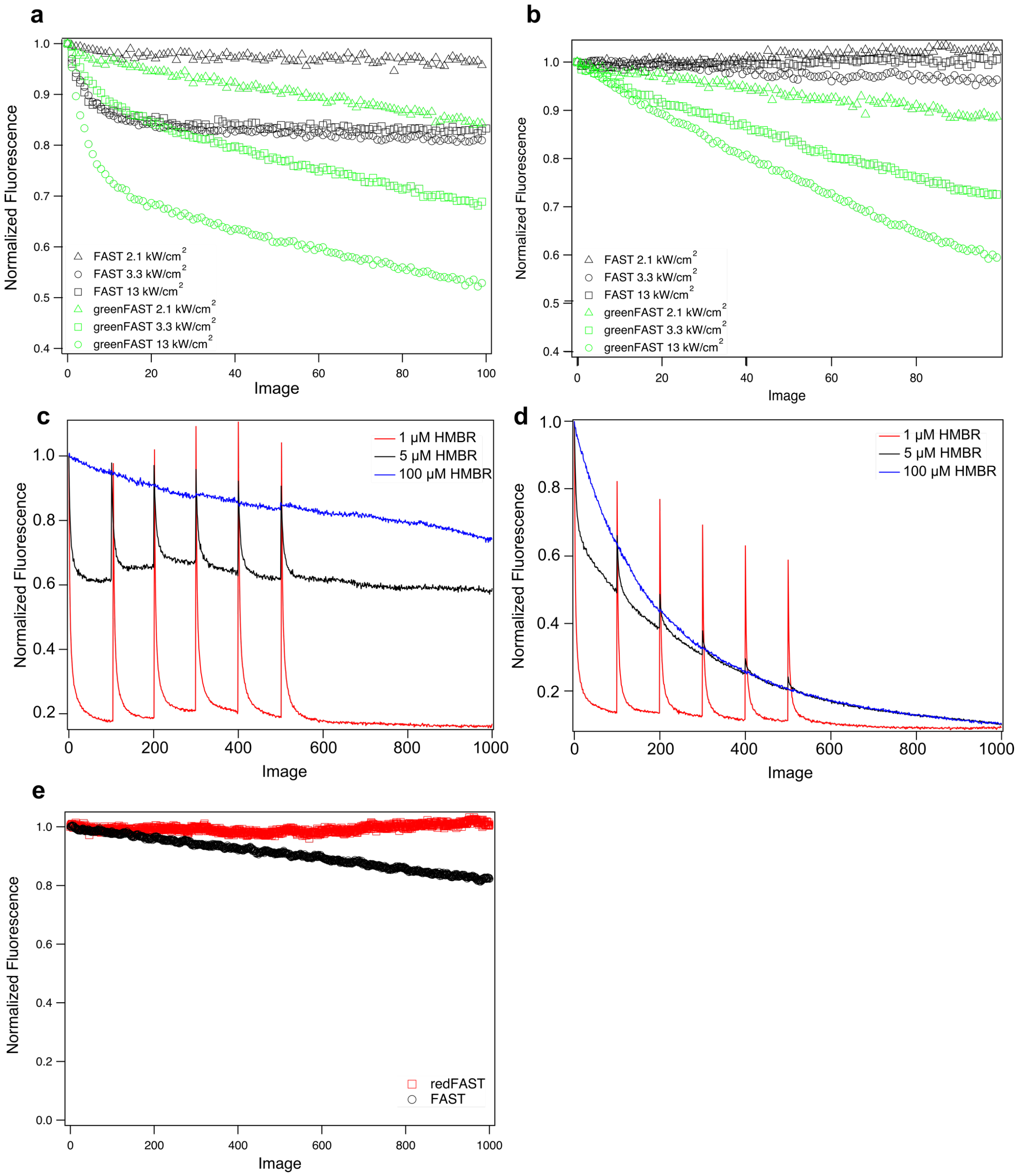


**Figure S3. Photostability measurements for greenFAST and redFAST. (a,b)** comparison of greenFAST and FAST photostability in different illumination conditions. Both were expressed as H2B fusions in HEK293T and imaged with 10 μM HMBR. Images taken every a) 1 s and b) 10 s with 1.27 μs pixel dwell, excitation with 488 nm laser. **(c,d)** Comparison of c) FAST and d) greenFAST photostability as a function of fluorogen concentration at 13 kW/cm$^2$ for 488 nm laser, 1.27 μs pixel dwell, images taken every 1s. 100 images were acquired followed by 60 s in the dark before acquisition was restarted. **(e)** redFAST and FAST expressed in the cytosol in HEK293T cells labeled with 10 μM HBR-3,5DOM were illuminated with 4.9 kW/cm$^2$ for 541 nm laser, 1.27 μs pixel dwell, images taken every 1 s. See **SI Text 1** for discussion of the results.

**Supplementary legends**

**Movie S1.** Zebrafish embryos were injected with redFAST-zGem(1-100)-P2A-greenFAST-zCdt1(1-190) mRNA at one-cell stage, and time-lapse imaging was performed starting from 256-cell stage on embryos incubated with 5 μM HMBR and 5 μM HBR-3,5DOM. Scale bar is 20 μm. Images every 5 minutes.

**Movie S2.** Whole zebrafish embryo imaging. Zebrafish embryos were injected with redFAST-zGem(1-100)-P2A-greenFAST-zCdt1(1-190) mRNA at one-cell stage, and time-lapse imaging was performed starting from 256-cell stage on embryos incubated with 5 μM HMBR and 5 μM HBR-3,5DOM. Scale bar is 100 μm.

**Movie S3.** Detection of FKBP-FKBP homodimer encoded by redFAST followed by detection of FRB-FKBP detected by greenFAST using time-lapse imaging in HEK293T cells labeled with 5 μM HMBR and 10 μM HBR-3,5DOM. First, the cells were treated with 100 nM AP1510 (imaged every 5 mins), and then at t = 140 min AP1510 was removed and 1 μM rapamycin was added (imaged every 30 s). Scale bar 20 μm.

**SI Text 1:**

Hybrid systems have been long proposed as a way to combine the advantages of organic fluorophores with those of proteins, namely being able to genetically target tags to particular cell types and as fusions with specific proteins. High photostability and photon budget are hallmarks of many organic fluorophores, which must be balanced with off-target labeling and high background signal. We have previously observed that while FAST systems do undergo photodestruction, they benefit from chromophore renewal due to the reversibility of the protein:fluorogen interaction (1). Furthermore, given fluorophores can differ in brightness when conjugated with either Halotag or SNAP-tag, indicating that even in the case of these highly promiscuous self-labeling tags, the protein tag portion may not be entirely innocent (2).

Here, we observe also that the photostability of FAST:HMBR and FAST:HBR-3,5DOM differ from greenFAST and redFAST, despite both new systems utilizing the same fluorogens. A total of 5 mutations confer nearly complete selectivity for HBR-3,5DOM to redFAST, while forming a complex with largely equivalent photophysical characteristics (absorptivity, fluorescence quantum yield, fluorescence lifetime). However, Figure S3e shows that these 5 mutations also confer greater resistance to photobleaching. In contrast, the 3 mutations introduced into greenFAST induce a dramatic difference in photostability. This effect was investigated at multiple illumination intensities and acquisition rates (Figure S3a-b), as well as at multiple fluorogen concentrations (Figure S3c-d). Both FAST and greenFAST exhibit partially reversible, biphasic behavior as has been observed previously due to photoisomerization of the chromophore and destruction of one or both components of the fluorescent complex (1). The data reported here can be described by a three-state model based on the previously published models (1) comprised of (irreversibly) photodamaged protein, free protein, and (reversible) protein:fluorogen complex. At high fluorogen concentrations the contribution of the reversible component is abrogated as protein is re-bound instantaneously and the irreversible photobleaching process dominates. Given the nature of the mutations introduced in greenFAST (G21E, P68T, G77R), it is not immediately obvious why greenFAST might be more sensitive to photodamage, but it is reasonable to hypothesize that this might be due to altered folding around the binding pocket, perhaps allowing greater solvent access. Nevertheless, imaging conditions can be chosen to optimize the signal — namely lower light intensities and lower total acquisitions. The photophysics of this class of fluorogen and the effect of the protein tag on its functional behavior are currently being investigated in detail.

### Table S1. Clones isolated from the green selection

| Clones | Number of appearances | Mutations | $K_D$ for HMBR (μM) | $K_D$ for HBR-3,5DOM (μM) |
|---|---|---|---|---|
| 1 | 7 | G21E, P68T, G77R | 0.09 | 16.2 |
| 2 | 1 | F62L, P68S, T70K, Y76F, K80N | 0.05 | 7.2 |
| 3 | 1 | P68T, T70K | n.d. | n.d. |
| 4 | 1 | S8R, F62L, P68H, T70P, N87D | n.d. | n.d. |
| 6 | 4 | P68T, F75L, E93D | 0.13 | 13.0 |
| 7 | 1 | G35S, D36G, S72T, E93D, V107M | 0.08 | 3.3 |
| 12 | 1 | P68T, T70R | n.d. | n.d. |
| 21 | 2 | Q41R, E93D, V107M | 0.10 | 4.4 |
| 24 | 1 | P68T, T70K, E93V, G115S | n.d. | n.d. |

n.d. not determined

**Table S2. Clones isolated from the red selection**

| Clones | Number of appearances | Mutations | $K_D$ for HMBR (μM) | $K_D$ for HBR-3,5DOM (μM) |
|---|---|---|---|---|
| 1 | 3 | K17R, D19G, F28L, A30T, E46Q, K60R | 0.92 | 0.51 |
| 2 | 7 | A30V, R52S, K60R, V83A, K111R, S117C, Y118F | 1.60 | 0.6 |
| 4 | 2 | G21R, F28L, E46Q | 0.98 | 0.78 |
| 5 | 1 | F28L, E46Q, S117R | 1.47 | 0.88 |
| 6 | 1 | L33F, Q41H, E46Q, K111N | 1.72 | 1.0 |
| 7 | 1 | R52A, K80M, S99I | 1.45 | 1.0 |
| 10 | 6 | R52A, E81V, S99N | 1.75 | 1.0 |
| 17 | 1 | D20H, F28I, E46Q | 1.1 | 0.84 |

### Table S3. Rationally designed clones

| Plasmid | Mutations | $K_D$ for HMBR (μM) | $K_D$ for HBR-3,5DOM (μM) |
|---|---|---|---|
| 302 | green clone 6 V107M | 0.05 | 16.4 |
| 303 | green clone 21 P68T | 0.13 | 15.6 |
| 304 | green clone 21 P68T T70K | 0.10 | 7.4 |
| 305 | green clone 21 V122I | 0.55 | 25.5 |
| 306 | red clone 7 I99N | 1.33 | 0.7 |
| 307 | red clone 10 F28L | 9.4 | 6.4 |
| 308 | red clone 10 F28L E46Q | 12 | 1.2 |

### Table S4. Average fluorescence lifetime determination of FAST:fluorogen complexes

| | | monoexponential fit | biexponential fit | |
|---|---|---|---|---|
| **protein** | **fluorogen** | **$\tau$ (ns)** | **$\tau_1$ (ns)** | **$\tau_2$ (ns)** |
| iFAST | HMBR | 1.50 ± 0.02 | 1.7 ± 0.02 | 0.7 ± 0.07 |
| iFAST | HBR-3,5DOM | 2.62 ± 0.06 | 2.77 ± 0.05 | 0.5 ± 0.2 |
| greenFAST | HMBR | 1.11 ± 0.01 | 1.18 ± 0.09 | 0.4 ± 0.4 |
| greenFAST | 5:10* | 1.10 ± 0.01 | 1.22 ± 0.09 | 0.6 ± 0.4 |
| redFAST | HBR-3,5DOM | 2.42 ± 0.03 | 2.48 ± 0.07 | 0.4 ± 0.5 |
| redFAST | 5:10* | 2.39 ± 0.05 | 2.46 ± 0.03 | 0.13 ± 0.09 |

* in presence of 5 μM HMBR + 10 μM HBR-3,5DOM

See Table S5-S10 for individual fit results.

### Table S5. Fluorescence lifetime determination of iFAST:HMBR.

| | monoexponential fit | | | biexponential fit | | | | |
|---|---|---|---|---|---|---|---|---|
| | A | $\tau$ (ns) | $\chi^2$ | A1 | $\tau_1$ (ns) | A2 | $\tau_2$ (ns) | $\chi^2$ |
| Cell1 | 10578.46 | 1.475 | 4.472 | 7092.61 | 1.716 | 4934.4 | 0.729 | 0.973 |
| Cell2 | 14741.84 | 1.509 | 4.826 | 10437.04 | 1.718 | 6104.15 | 0.743 | 0.987 |
| Cell3 | 13313.04 | 1.471 | 5.762 | 8398.85 | 1.742 | 6707.23 | 0.772 | 1.029 |
| Cell4 | 4073.04 | 1.509 | 1.935 | 3063.62 | 1.689 | 1523.5 | 0.679 | 0.975 |
| Cell5 | 7365.25 | 1.519 | 2.77 | 5432.43 | 1.711 | 2841.34 | 0.709 | 0.976 |
| Cell6 | 14109.8 | 1.485 | 4.934 | 9950.92 | 1.695 | 5976.06 | 0.72 | 0.994 |
| Cell7 | 3942.55 | 1.511 | 1.9 | 3024.85 | 1.682 | 1433.01 | 0.65 | 0.961 |
| Cell8 | 4792.98 | 1.512 | 2.012 | 3518.86 | 1.703 | 1838.22 | 0.725 | 0.922 |
| Cell9 | 7427.48 | 1.513 | 2.724 | 5139.95 | 1.731 | 3103.13 | 0.794 | 1.01 |
| Cell10 | 11558.12 | 1.468 | 5.301 | 7410.5 | 1.732 | 5775.46 | 0.747 | 1.069 |
| Cell11 | 4528.78 | 1.504 | 1.969 | 3358.88 | 1.689 | 1723.41 | 0.7 | 0.911 |
| Cell12 | 5223.34 | 1.479 | 2.603 | 3818.87 | 1.676 | 2211.79 | 0.643 | 0.924 |
| Cell13 | 11762.86 | 1.498 | 4.163 | 8864.49 | 1.677 | 4537.6 | 0.645 | 0.961 |
| Cell14 | 6579.2 | 1.487 | 3.014 | 4726.88 | 1.693 | 2805.27 | 0.67 | 0.934 |
| Cell15 | 8212.45 | 1.515 | 2.748 | 6425.35 | 1.673 | 2820.98 | 0.645 | 0.988 |
| Cell16 | 5028.68 | 1.508 | 2.009 | 3516.45 | 1.717 | 2039.22 | 0.798 | 0.954 |
| Cell17 | 6203.44 | 1.492 | 2.981 | 4239.95 | 1.726 | 2806.52 | 0.729 | 1.026 |
| Cell18 | 4904.13 | 1.494 | 2.69 | 3928.75 | 1.642 | 1899.4 | 0.492 | 1.131 |
| Cell19 | 9044.41 | 1.478 | 3.958 | 5851.78 | 1.736 | 4399.14 | 0.763 | 0.911 |
| Cell20 | 12247.06 | 1.486 | 4.79 | 8476.43 | 1.707 | 5404.3 | 0.725 | 1.055 |

### Table S6. Fluorescence lifetime determination of greenFAST:HMBR

| | monoexponential fit | | | biexponential fit | | | | |
|---|---|---|---|---|---|---|---|---|
| | A | $\tau$ (ns) | $\chi^2$ | A1 | $\tau_1$ (ns) | A2 | $\tau_2$ (ns) | $\chi^2$ |
| Cell1 | 27370.09 | 1.113 | 2.057 | 26939.49 | 1.121 | 26696.94 | 0.033 | 1.664 |
| Cell2 | 14098.96 | 1.128 | 1.395 | 11391.21 | 1.198 | 3150.07 | 0.726 | 1.092 |
| Cell3 | 17045.11 | 1.112 | 1.706 | 16735.95 | 1.121 | 14765.24 | 0.038 | 1.406 |
| Cell4 | 9503.22 | 1.132 | 1.227 | 5862.89 | 1.259 | 4022.13 | 0.876 | 1.022 |
| Cell5 | 13197.7 | 1.129 | 1.293 | 12972.02 | 1.141 | 1302.8 | 0.163 | 1.15 |
| Cell6 | 13836.96 | 1.129 | 1.337 | 11161.69 | 1.205 | 3318.65 | 0.729 | 0.999 |
| Cell7 | 18519.36 | 1.112 | 1.369 | 7338.28 | 1.306 | 11600.95 | 0.948 | 0.996 |
| Cell8 | 21487.22 | 1.115 | 1.544 | 21224.2 | 1.122 | 2826.47 | 0.117 | 1.333 |
| Cell9 | 16927.76 | 1.11 | 1.393 | 16522.99 | 1.115 | 3091.94 | 0.08 | 1.248 |
| Cell10 | 26328.43 | 1.086 | 2.526 | 17419.74 | 1.216 | 10552.16 | 0.736 | 1.068 |
| Cell11 | 17946.31 | 1.103 | 1.725 | 17647.59 | 1.11 | 18925.53 | 0.031 | 1.485 |
| Cell12 | 4859.09 | 1.121 | 1.133 | 4710.63 | 1.129 | 11053.38 | 0.014 | 1.036 |
| Cell13 | 6036.19 | 1.122 | 1.128 | 1799.51 | 1.417 | 4442.82 | 0.954 | 0.888 |
| Cell14 | 9214.84 | 1.078 | 1.508 | 9018.14 | 1.088 | 7124.8 | 0.044 | 1.309 |
| Cell15 | 4349.51 | 1.107 | 0.9 | 3387.54 | 1.19 | 1165.22 | 0.741 | 0.791 |

**Table S7. Fluorescence lifetime determination of greenFAST in presence of both fluorogens (5 μM HMBR + 10 μM HBR-3,5DOM)**

| | monoexponential fit | | | biexponential fit | | | | |
|---|---|---|---|---|---|---|---|---|
| | **A** | **$\tau$ (ns)** | **$\chi^2$** | **A1** | **$\tau_1$ (ns)** | **A2** | **$\tau_2$ (ns)** | **$\chi^2$** |
| Cell1 | 11173.58 | 1.078 | 1.303 | 7705.49 | 1.194 | 4133.85 | 0.725 | 0.768 |
| Cell2 | 6930.13 | 1.105 | 1.248 | 6203.97 | 1.156 | 1177.52 | 0.504 | 0.987 |
| Cell3 | 18878.47 | 1.107 | 1.7 | 18509.18 | 1.113 | 33085.25 | 0.019 | 1.472 |
| Cell4 | 5446.24 | 1.107 | 0.847 | 5236.14 | 1.112 | 72432.55 | 0.003 | 0.804 |
| Cell5 | 10142.73 | 1.076 | 1.76 | 9783.5 | 1.093 | 6448.44 | 0.064 | 1.369 |
| Cell6 | 16119.46 | 1.104 | 1.577 | 8451.9 | 1.274 | 8452.32 | 0.859 | 0.958 |
| Cell7 | 15506.13 | 1.113 | 1.43 | 15037.19 | 1.116 | 9696.21 | 0.02 | 1.355 |
| Cell8 | 16499.81 | 1.089 | 1.733 | 9112.22 | 1.252 | 8263.47 | 0.828 | 0.967 |
| Cell9 | 5101.52 | 1.124 | 1.029 | 5019.57 | 1.135 | 582.18 | 0.14 | 0.975 |
| Cell10 | 7193.56 | 1.103 | 1.143 | 3666.89 | 1.287 | 3891.32 | 0.855 | 0.841 |
| Cell11 | 15727.38 | 1.112 | 1.359 | 7656.04 | 1.286 | 8733.3 | 0.902 | 0.933 |
| Cell12 | 24572.01 | 1.097 | 1.928 | 13650.78 | 1.255 | 12148.35 | 0.838 | 0.971 |
| Cell13 | 12991.7 | 1.106 | 1.388 | 4386.7 | 1.353 | 9150.58 | 0.944 | 0.978 |
| Cell14 | 12118.07 | 1.109 | 1.379 | 3783.49 | 1.383 | 8864.49 | 0.947 | 0.94 |
| Cell15 | 13471.46 | 1.095 | 1.386 | 8461.83 | 1.227 | 5686.63 | 0.808 | 0.898 |

### Table S8. Fluorescence lifetime determination of iFAST:HBR-3,5DOM

| | monoexponential fit | | | biexponential fit | | | | |
|---|---|---|---|---|---|---|---|---|
| | **A** | **$\tau$ (ns)** | **$\chi^2$** | **A1** | **$\tau_1$ (ns)** | **A2** | **$\tau_2$ (ns)** | **$\chi^2$** |
| Cell1 | 1965.42 | 2.586 | 1.947 | 1736.53 | 2.77 | 704.13 | 0.457 | 1.078 |
| Cell2 | 2639.56 | 2.642 | 1.782 | 2412.05 | 2.778 | 662.7 | 0.496 | 1.172 |
| Cell3 | 2201.31 | 2.596 | 1.918 | 1950.74 | 2.777 | 723.07 | 0.49 | 1.072 |
| Cell4 | 4469.52 | 2.609 | 2.381 | 4010.16 | 2.768 | 1259.21 | 0.525 | 1.049 |
| Cell5 | 4425.4 | 2.678 | 1.637 | 3846.89 | 2.872 | 930.57 | 1.001 | 1.022 |
| Cell6 | 2653.59 | 2.611 | 2.162 | 2373.2 | 2.776 | 926.74 | 0.425 | 1.077 |
| Cell7 | 7151.08 | 2.595 | 3.627 | 6395.55 | 2.761 | 2237.1 | 0.484 | 1.083 |
| Cell8 | 6423.93 | 2.612 | 3.102 | 5758.78 | 2.773 | 1854.14 | 0.517 | 1.081 |
| Cell9 | 2456.01 | 2.448 | 3.813 | 2017.73 | 2.743 | 1425.48 | 0.434 | 1.078 |
| Cell10 | 1910.88 | 2.676 | 1.237 | 1880.57 | 2.681 | 1928.43 | 0.026 | 1.13 |
| Cell11 | 2501.35 | 2.645 | 1.416 | 2325.02 | 2.757 | 532.52 | 0.477 | 1.02 |
| Cell12 | 1688.3 | 2.666 | 1.257 | 1532.78 | 2.813 | 346.11 | 0.671 | 0.988 |
| Cell13 | 1996.75 | 2.677 | 1.237 | 1837.91 | 2.803 | 374.38 | 0.635 | 0.969 |
| Cell14 | 1747.61 | 2.678 | 1.242 | 1631.1 | 2.789 | 316.94 | 0.537 | 1.029 |
| Cell15 | 2607.02 | 2.647 | 1.495 | 2558.18 | 2.668 | 1694.42 | 0.072 | 1.304 |

### Table S9. Fluorescence lifetime determination of redFAST:HBR-3,5DOM

| | monoexponential fit | | | biexponential fit | | | | |
|---|---|---|---|---|---|---|---|---|
| | A | $\tau$ (ns) | $\chi^2$ | A1 | $\tau_1$ (ns) | A2 | $\tau_2$ (ns) | $\chi^2$ |
| Cell1 | 1814.86 | 2.396 | 1.208 | 1713.03 | 2.478 | 331.12 | 0.418 | 0.997 |
| Cell2 | 2414.63 | 2.455 | 1.072 | 2281.37 | 2.53 | 265.37 | 0.76 | 0.97 |
| Cell3 | 2063.17 | 2.431 | 1.132 | 2042.45 | 2.437 | 2392.74 | 0.032 | 1.059 |
| Cell4 | 2957.77 | 2.445 | 1.167 | 2919.34 | 2.458 | 1086.28 | 0.055 | 1.097 |
| Cell5 | 828.31 | 2.347 | 1.208 | 768.06 | 2.446 | 399.47 | 0.19 | 0.887 |
| Cell6 | 891.13 | 2.45 | 0.898 | 744.72 | 2.625 | 183.3 | 1.391 | 0.853 |
| Cell7 | 765.46 | 2.41 | 1.038 | 787.37 | 2.413 | -1518.57 | 0.031 | 1.021 |
| Cell8 | 979.48 | 2.439 | 0.918 | 911.8 | 2.531 | 124.54 | 0.824 | 0.867 |
| Cell9 | 515.49 | 2.374 | 0.896 | 515.73 | 2.374 | -155.36 | 0 | 0.898 |
| Cell10 | 1863.2 | 2.455 | 1.026 | 1522.06 | 2.652 | 425.65 | 1.38 | 0.911 |
| Cell11 | 3337.64 | 2.454 | 1.179 | 3300.5 | 2.458 | 3716.89 | 0.012 | 1.075 |
| Cell12 | 2921.11 | 2.445 | 1.135 | 2891.01 | 2.449 | 2692.52 | 0.015 | 1.081 |
| Cell13 | 1938.98 | 2.415 | 1.227 | 1817.72 | 2.507 | 374.3 | 0.442 | 0.967 |
| Cell14 | 1436.13 | 2.424 | 1.01 | 1412.72 | 2.437 | 699.3 | 0.04 | 0.958 |
| Cell15 | 2346.33 | 2.451 | 0.989 | 2344.62 | 2.452 | 1060.38 | 0 | 0.991 |
| Cell16 | 1432.24 | 2.416 | 1.124 | 1325.28 | 2.524 | 264.85 | 0.557 | 0.939 |
| Cell17 | 1479.55 | 2.41 | 1.053 | 1457.19 | 2.431 | 811.07 | 0.056 | 0.965 |

**Table S10. Fluorescence lifetime determination of redFAST in presence of both fluorogens (5 µM HMBR + 10 µM HBR-3,5DOM)**

| | monoexponential fit | | | biexponential fit | | | | |
|---|---|---|---|---|---|---|---|---|
| | **A** | **$\tau$ (ns)** | **$\chi^2$** | **A1** | **$\tau_1$ (ns)** | **A2** | **$\tau_2$ (ns)** | **$\chi^2$** |
| Cell1 | 1428.92 | 2.262 | 3.688 | 1254.12 | 2.423 | 2067.49 | 0.119 | 1.04 |
| Cell2 | 3816.95 | 2.378 | 2.664 | 3596.76 | 2.445 | 2731.71 | 0.108 | 1.057 |
| Cell3 | 3905.82 | 2.434 | 1.486 | 3719.87 | 2.504 | 678.7 | 0.388 | 1.08 |
| Cell4 | 7004.25 | 2.43 | 1.511 | 6923.67 | 2.444 | 9689.39 | 0.028 | 1.192 |
| Cell5 | 6466.97 | 2.429 | 2.127 | 6160.42 | 2.496 | 1506.33 | 0.284 | 1.172 |
| Cell6 | 2483.86 | 2.443 | 1.648 | 2365.88 | 2.507 | 1096.33 | 0.147 | 1.049 |
| Cell7 | 6634.34 | 2.43 | 1.489 | 6571.84 | 2.441 | 8457.66 | 0.029 | 1.265 |
| Cell8 | 3026.95 | 2.38 | 1.853 | 2868.4 | 2.449 | 1241.79 | 0.171 | 1.088 |
| Cell9 | 3683.57 | 2.398 | 1.544 | 3551.47 | 2.445 | 1203.28 | 0.151 | 1.038 |
| Cell10 | 4610.46 | 2.434 | 1.447 | 4561.21 | 2.446 | 6194.79 | 0.029 | 1.23 |
| Cell11 | 4329.77 | 2.341 | 3.864 | 4045.72 | 2.426 | 3852.1 | 0.104 | 1.171 |
| Cell12 | 4008.91 | 2.274 | 8.293 | 3577.52 | 2.406 | 6120.45 | 0.101 | 1.323 |
| Cell13 | 13575.71 | 2.405 | 7.021 | 12842.39 | 2.463 | 10664.73 | 0.094 | 1.375 |
| Cell14 | 5577.29 | 2.402 | 5.794 | 5189.37 | 2.489 | 6302.6 | 0.09 | 1.215 |
| Cell15 | 5242.42 | 2.368 | 5.231 | 4875 | 2.46 | 4898.98 | 0.113 | 1.203 |

**Table S11. Sequence of oligonucleotides used in this study**

| Name | Sequence |
|---|---|
| ag175 | gcagcggcggagggggatccatggagcatgttgcctttggc |
| ag182 | ggatccccctccgccgctgccgcctcctccggagacctgctttgagattcgtcgg |
| ag184 | ggatccccctccgccgctgccgcctcctccggattcttccagttttagaagctccacatc |
| ag189 | gtggtgctcgagctattactacacccgttttataaagacccaatagc |
| ag195 | gcctgtgatgtctccctgagcagcattgtac |
| ag196 | tacaatgctgctcagggagacatcacaggc |
| ag216 | ttcgtagctagcatggagcatgttgcctttg |
| ag217 | ttgttcggatcccacccgtttcacaaagac |
| ag224 | atggctagcgaaaacctgtattttcagggcatggagcatgttgcctttggc |
| ag311 | aaagcttatttctgaagaggacttgtaataggcggccgcgactctagatcataatc |
| ag313 | ctcaccttgctcctgccgagaaagtatcca |
| ag314 | tggatactttctcggcaggagcaaggtgag |
| ag321 | gccctgaaaatacaggttttcgctagc |
| ag322 | taatagctcgagcaccaccaccac |
| ag347 | taataggcggccgcgactctag |
| ag354 | gtggtggtggtgctcgagctattacacccgtttcacaaagacccaatag |
| ag356 | gtcctcttcagaaataagcttttgttcggatcccacccgtttcacaaagacccaatag |
| ag357 | ccggactcagatctgccaccatggagcatgttgcctttggcag |
| ag358 | ggtggcagatctgagtccggtag |
| ag420 | ccaaccaaggtcaagatgcacatgaagaaag |
| ag421 | ctttcttcatgtgcatcttgaccttggttgg |
| ag422 | caaggatgttgcaactggaacggattctc |
| ag423 | gagaatccgttccagttgcaacatccttg |
| ag424 | gatgttgcaactggaaaggattctcccgag |
| ag425 | ctcgggagaatcctttccagttgcaacatc |
| ag426 | gaatggatgataccgacaaacaggggaccaaccaag |
| ag427 | cttggttggtcccctgtttgtcggtatcatccattc |
| ag428 | gatgggttggccttaggcgcaattcagctc |
| ag429 | gagctgaattgcgcctaaggccaacccatc |
| ag472 | ctagagtcgcggccgcctattaggaaagggctttcttcatgtgcac |
| ag491 | ggaggcggatctgccaccatggagcatgttgcctttggcag |
| ag492 | catggtggcagatccgcctcc |
| ag527 | gccaaaggcaacatgctccatgaattccaagtcctcttcagaaataagcttttgttc |
| ag528 | atggagcatgttgcctttgg |
| ag530 | cacccgtttcacaaagaccc |
| ag532 | gctgaagcaggctggagacgtggaggagaaccctggacctatggagcatgttgcctttgg |
| ag533 | gggtctttgtgaaacgggtgggatatccatcacactggcgg |
| ag534 | ctagagtcgcggccgcctattacagcgcctttctccgtttttc |
| ag550 | tgctgaagcaggctggagacgtggaggagaaccctggacctgtgagcaagggcgaggagg |
| ag554 | catcaagtccaagggcaaggactccgccggcggcggctccatggagcatgttgcctttggc |
| ag555 | gagtccttgcccttggacttgatg |
| ag598 | tacagcatgctgccgagc |

| | |
|---|---|
| ag599 | cctgcttcagcaggctgaagttagtagctccgcttcctatagtgtcctgatcctgggctg |
| ag675 | gctcggcagcatgctgtacacccgtttcacaaagaccc |
| ag677 | ctaccggactcagatctgccaccatgggcgtggccgacttgatcaagaagttcgagtcca |
| ag678 | ctccatgaccggtggatccccctcctccttggagatggactcgaacttcttgatcaagtc |
| ag679 | ggagggggatccaccggtcatggagcatgttgcctttgg |
| ag795 | ctaccggactcagatctgccaccatggtgtcccggcaagaagaag |
| ag796 | ccaaaggcaacatgctccatgtctggtttaatcacactcatggtgg |
| Kan-F | gcatcaaccaaaccgttattcattcgtg |
| Kan-R | cacgaatgaataacggtttggttgatgc |

### Table S12. Table of plasmids used in this study

| Plasmid code | Expression host | open reading frame |
|---|---|---|
| pAG261 | *E. coli* | green clone 1 = greenFAST |
| pAG262 | *E. coli* | green clone 2 |
| pAG263 | *E. coli* | green clone 6 |
| pAG264 | *E. coli* | green clone 7 |
| pAG265 | *E. coli* | green clone 21 |
| pAG270 | *E. coli* | red clone 1 |
| pAG271 | *E. coli* | red clone 2 |
| pAG272 | *E. coli* | red clone 4 |
| pAG273 | *E. coli* | red clone 5 |
| pAG274 | *E. coli* | red clone 6 |
| pAG275 | *E. coli* | red clone 7 |
| pAG276 | *E. coli* | red clone 10 |
| pAG277 | *E. coli* | red clone 17 |
| pAG302 | *E. coli* | green clone 6 V107M |
| pAG303 | *E. coli* | green clone 21 P68T |
| pAG304 | *E. coli* | green clone 21 P68T T70K |
| pAG305 | *E. coli* | green clone 21 V122I |
| pAG306 | *E. coli* | red clone 7 I99N |
| pAG307 | *E. coli* | red clone 10 F28L |
| pAG308 | *E. coli* | red clone 10 F28L E46Q = redFAST |
| pAG361 | Mammalian | lyn11-greenFAST |
| pAG362 | Yeast | redFAST |
| pAG364 | Mammalian | greenFAST |
| pAG365 | Mammalian | redFAST |
| pAG369 | Mammalian | lyn11-redFAST |
| pAG372 | Mammalian | mito-greenFAST |
| pAG373 | Mammalian | mito-redFAST |
| pAG374 | Mammalian | H2B-greenFAST |
| pAG375 | Mammalian | H2B-redFAST |
| pAG460 | Mammalian | FRB-N-greenFAST |
| pAG461 | Mammalian | FRB-N-redFAST |
| pAG462 | Mammalian | FKBP-N-greenFAST |
| pAG463 | Mammalian | FKBP-N-redFAST |
| pAG469 | Mammalian | LifeAct-redFAST |
| pAG477 | Mammalian | redFAST-Cdt(30-120)-P2A-greenFAST-Gem(1-120) |
| pAG551 | Mammalian | MAP4-greenFAST |
| pAG552 | Mammalian | MAP4-redFAST |
| pAG646 | Yeast | green clone 6 V107M |
| pAG647 | Yeast | green clone 21 P68T |
| #1113 | Vertebrate | pT2iC6-LifeAct-greenFAST |
| #1135 | Vertebrate | redFAST-zGem(1-100)-P2A-greenFAST-zCdt1(1-190) |

### Materials and Methods

#### General

Synthetic oligonucleotides used for cloning were purchased from Integrated DNA Technology. The sequences of oligonucleotides used in this study are provided in **Table S11**. PCR reactions were performed with Q5 polymerase (New England Biolabs) in the buffer provided. PCR products were purified using QIAquick PCR purification kit (Qiagen). The products of restriction enzyme digests were purified by preparative gel electrophoresis followed by QIAquick gel extraction kit (Qiagen). Restriction endonucleases, T4 ligase, Phusion polymerase, Taq ligase, and Taq exonuclease were purchased from New England Biolabs and used with accompanying buffers and according to manufacturer protocols. Isothermal assemblies (Gibson assembly) were performed using homemade mix prepared according to previously described protocols (3). Small-scale isolation of plasmid DNA was done using QIAprep miniprep kit (Qiagen) from 2 mL of overnight culture. Large-scale isolation of plasmid DNA was done using the QIAprep maxiprep kit (Qiagen) from 150 mL of overnight culture. All plasmid sequences were confirmed by Sanger sequencing with appropriate sequencing primers (GATC Biotech). Please see **Table S12** for a list of all plasmids used in this study. The preparation of HMBR (4-hydroxy-3-methylbenzylidene rhodanine) and HBR-3,5DOM (4-hydroxy-3,5-dimethoxybenzylidene rhodanine) were previously described[2,3]. HMBR and HBR-3,5DOM are available from The Twinkle Factory under the name $^{TF}$Lime and $^{TF}$Coral (thetwinklefactory.com).

#### Yeast Display

*Library construction.* The yeast display libraries of YFAST were constructed by error-prone PCR using the Genemorph II kit (Agilent) using primers ag216/ag217. The error rate of the PCR was varied by using either 1 or 10 ng template gene. The PCR reactions were mixed to achieve a mutation rate of 3.8 nt/gene and cloned into pCTCON2 using NheI and BamHI restriction sites. Large scale transformation into DH10B was performed, yielding $10^6$ transformants. The DNA was maxiprepped and transformed into yeast strain EBY100 using a large-scale, high-efficiency protocol (4) to yield $2 \times 10^6$ transformants.

*Selection.* The library ($1.5 \times 10^9$ cells) was grown overnight at 30 °C in 1 L SD (20 g/L dextrose, 6.7 g/L yeast nitrogen base, 1.92 g/L yeast synthetic dropout without tryptophan, 7.44 g/L $NaH_2PO_4$, 10.2 g/L $Na_2HPO_4$-$7H_2O$, 1% penicillin-streptomycin 10,000 U/mL). The following morning the culture was diluted to $OD_{600nm}$ in 1 L SD and grown at 30 °C until the $OD_{600nm}$ was between 2 and 5 at which point $5 \times 10^6$ cells were pelleted and resuspended in SG (20 g/L galactose, 2 g/L dextrose, 6.7 g/L yeast nitrogen base, 1.92 g/L yeast synthetic dropout without tryptophan, 7.44 g/L $NaH_2PO_4$, 10.2 g/L $Na_2HPO_4$-$7H_2O$, 1% penicillin-streptomycin 10,000 U/mL). The cultures were then grown for 36 h at 23 °C. $1.2 \times 10^9$ cells were pelleted by centrifugation (2500 × *g*, 3 min), washed once with 10 mL DPBS + BSA (1 g/L) and incubated for 30 min at room temperature in 480 μL of a 1:250 dilution of chicken anti-c-myc IgY (Life Technologies) in DPBS-BSA. Cells were then centrifuged and washed with DPBS-BSA and incubated for 20 min on ice with a 1:150 dilution of secondary goat anti-chicken coupled to Alexa-Fluor 647. After centrifugation and washing with DPBS-BSA, the cells were resuspended in 10 mL DPBS-BSA supplemented with either 5 μM HMBR + 5 μM HBR-3,5DOM, or 1 μM HMBR + 10 μM HBR-3, 5DOM. The cells were sorted on a MoFlo Astrios (Beckman Coulter) equipped with 488 nm, 561 nm, and 640 nm laser. Sorted cells were collected in SD, grown over night at 30 °C, and plated on SD agar plates. The plates were incubated for 3 days at 30 °C and the resulting lawn was resuspended in SD supplemented with 20% glycerol. The resulting stock was either frozen or used directly for the next round as well as tested for viability by serial dilution and plating on SD plates. After 4-5 rounds of selection by FACS, 24 clones were screened by flow cytometry and their DNA was isolated using a miniprep kit (Qiagen), transformed into DH10B and re-isolated for sequencing.

**Cloning**

Selected clones were subcloned into a pET28a backbone for recombinant expression in *E. coli* by isothermal assembly using backbone fragments generated by PCR amplification of pAG87(5) using primers ag321/KanF and ag322/KanR and inserts amplified from the isolated yeast plasmids using primers ag354 and ag224.

Site-directed mutagenesis was carried out using isothermal assembly of two overlapping PCR fragments with mutations encoded on primers. The plasmid pAG302 was generated by introduction of the V107M mutation by amplification of pAG263 with primers ag420/KanR and ag421/KanF. The plasmid pAG303 was generated by introduction of the P68T mutation by amplification of pAG265 with primers ag422/KanR and ag423/KanF. The plasmid pAG304 was generated by introduction of the T70K mutation by amplification of pAG303 with primers ag423/KanR and ag424/KanF. The plasmid pAG305 was generated by introduction of the V122I mutation by amplification of pAG265 with primers ag189/KanR and ag322/KanF. The plasmid pAG306 was generated by introduction of the I99N mutation by amplification of pAG2275 with primers ag426/KanR and ag427/KanF. The plasmid pAG307 was generated by introduction of the F28L mutation by amplification of pAG276 with primers ag428/KanR and ag429/KanF. The plasmid pAG308 was generated by introduction of the E46Q mutation by amplification of pAG307 with primers ag195/KanF and ag196/KanR.

The plasmids pAG364 and pAG365 encoding greenFAST and redFAST were constructed by isothermal assembly from the plasmid pAG104(5) (ref) encoding FAST. The sequences for the inserts encoding greenFAST and redFAST were amplified by PCR from pAG261 and pAG308 using primers ag356/ag357. The backbone was amplified using ag358/ag313 and ag311/ag314.

The plasmids pAG361 and pAG369 encoding lyn11-greenFAST and lyn11-redFAST, respectively, were constructed by isothermal assembly from the plasmid pAG106(5) encoding lyn11-FAST. The sequences encoding for lyn11-greenFAST and lyn11-redFAST were amplified by PCR from pAG261 and pAG308 using primers ag554/ag356. The backbone of pAG106 was amplified using ag555/ag313.

The plasmids pAG372 and pAG373 were generated by digestion pAG364 and pAG365 with BglII and HindIII and insertion into pAG156(5), which encodes for mito-FAST.

The plasmids pAG374 and pAG375 encoding H2B-greenFAST and H2B-redFAST were constructed by isothermal assembly from the plasmid pAG109, which encodes for H2B-FAST. The sequences encoding for greenFAST and redFAST were amplified from pAG261 and pAG308 using primers ag491/ag356. The backbone was amplified using ag492/ag313.

The plasmids pAG551 and pAG552 encoding MAP4-greenFAST and MAP4-redFAST were constructed by isothermal assembly from pAG364 and pAG365 encoding greenFAST and redFAST. The sequence for MAP4 was amplified from plasmid ffDronpa-MAP4 (6) using primers ag795/ag796. The backbones were amplified using ag528/ag313 and ag358/ag313.

The plasmids pAG469 and #1113 encoding LifeAct-redFAST and LifeAct-greenFAST were constructed by isothermal assembly from pAG365 and pAG364, respectively. The backbone was amplified from pAG364 and pAG365 encoding CMV-greenFAST and CMV-redFAST using primers ag679/ag314 and ag358/ag313 and assembled with ag677 and ag678.

The plasmid pAG477 encoding redFAST-Cdt(30-120)-P2A-greenFAST-Gem(1-120) was constructed in multiple steps by isothermal assembly from pAG148(7). The sequence for redFAST was amplified from pAG308 using primers ag528/ag314. The backbone was amplified using ag527/ag313. The sequence for Cdt(30-120) was amplified from a synthesized fragment (Eurofins) using primers ag598/ag599. The backbone was amplified using primers ag550/ag314 and ag675/ag313 (pAG439). The sequences for greenFAST and Geminin(1-120) were amplified from pAG261 and a synthesized fragment (Eurofins) using primers ag532/ag530 and ag533/ag534. The backbone was amplified using primer ag599/ag313 and ag347/314.

The plasmids pAG460 and pAG461 encoding FRB-N-greenFAST and FRB-N-redFAST were generated by isothermal assembly from pAG149 encoding FRB-NFAST. The sequences encoding N-greenFAST and N-redFAST were amplified from pAG364 and pAG365 using ag175/ag472. The backbones were amplified using ag182/ag313 and ag347/314.

The plasmids pAG462 and pAG463 encoding FKBP-N-greenFAST and FKBP-N-redFAST were generated by isothermal assembly from pAG148 encoding FKBP-NFAST. The sequences encoding N-greenFAST and N-redFAST were amplified from pAG364 and pAG365 using ag175/ag472. The backbones were amplified using ag184/ag313 and ag347/314.

The plasmids pAG362, pAG646, and pAG647 were generated from the yeast display plasmid pCTCON2 by restriction enzyme cloning using NheI and BamHI. The inserts were amplified using ag216/ag217 from pAG308, pAG302, and pAG303.

**Flow Cytometry**

Flow cytometry was performed on a MACSQuant Analyzer equipped with 405 nm, 488 nm, and 561 nm lasers and eight filters and channels. To prepare samples for flow cytometry, small scale cultures were grown as for library expression (*vide supra*). Briefly, 3 mL of SD were inoculated with a single colony and grown overnight at 30 °C. The following day, the cultures were diluted to a final $OD_{600nm}$ 1 in 5 mL of SD and grown until doubled. These cultures were used to inoculate 5 mL of either SD (non-induced) or SG (induced) to an OD of 0.5 and the cultures were grown for 36 h at 23°C. $1 \times 10^8$ cells were pelleted and washed with 1× DPBS + BSA. Aliquots of each were labeled with chicken anti-myc antibody as for library preparation using a secondary goat anti-chicken coupled to Alexa-Fluor 488 to verify protein expression. Clones were finally resuspended in 1× DPBS + BSA supplemented with one of the following fluorogen conditions: 0 μM, 5 μM HMBR, 10 μM HBR-3,5DOM, 1 μM HMBR + 10 μM HBR-3,5DOM, 5 μM HMBR + 5 μM HBR-3,5DOM, 5 μM HMBR + 10 μM HBR-3,5DOM. Data were analyzed in R (3.6.2) using RStudio with opencyto, flowCore, and ggcyto packages.

**Protein Expression and purification**

Expression vectors were transformed in Rosetta (DE3) pLysS *E. coli* (New England Biolabs). Cells were grown at 37°C in LB medium complemented with 50 μg/ml kanamycin and 34 μg/ml chloramphenicol to $OD_{600nm}$ 0.6. Expression was induced for 4 h by adding isopropyl β-D-1-thiogalactopyranoside (IPTG) to a final concentration of 1 mM. Cells were harvested by centrifugation (4,000 × g for 20 min at 4°C) and frozen. The cell pellet was resuspended in 1× Tris-EDTA-sucrose (TES) buffer (8) and incubated for 1 hr. The lysate was then diluted by three using 0.25 × TES buffer and incubated for 45 min. Cellular fragments were removed by centrifugation (9200 × g for 1.5 h at 4°C). The supernatant was incubated overnight at 4°C under gentle agitation with Ni-NTA agarose beads in phosphate buffered saline (PBS) (sodium phosphate 50 mM, NaCl 150 mM, pH 7.4) complemented with 10 mM imidazole. Beads were washed with ~20 volumes of PBS containing 20 mM imidazole, and with ~5 volumes of PBS complemented with 40 mM imidazole. His-tagged proteins were eluted with ~5 volumes of PBS complemented with 0.5 M imidazole. The buffer was exchanged to PBS (50 mM phosphate, 150 mM NaCl, pH 7.4) using PD-10 desalting columns. Purity of the proteins was evaluated using SDS-PAGE electrophoresis stained with Coomassie blue.

**Physico-chemical Measurements**

Steady state UV-Vis absorption spectra were recorded using a Cary 300 UV-Vis spectrometer (Agilent Technologies), equipped with a Versa20 Peltier-based temperature-controlled cuvette chamber (Quantum Northwest) and fluorescence data were recorded using a LPS 220 spectrofluorometer (PTI, Monmouth Junction, NJ), equipped with a TLC50TM Legacy/PTI Peltier-based temperature-controlled cuvette chamber (Quantum Northwest).

Thermodynamic dissociation constants and quantum yield measurements for HMBR or HBR-3,5DOM were determined as previously described using either FAST:HMBR or FAST:HBR-3,5DOM as a reference (5, 9, 10). Thermodynamic dissociation constants were determined with a Spark 10M plate reader (Tecan) and fit in Prism 6 to a one-site specific binding model. Quantum yield measurments were determined by reciprocal dilution with protein solution so as to keep the protein concentration constant at 40 μM and varying the concentration only of the protein:fluorogen complex. Absorption coefficients were determined by forward titration of fluorogen into a 40 μM protein solution using FAST as standard for the concentration of the fluorogen solution.

**Mammalian cell culture**

HEK 293T and COS-7 cells were cultured in Dulbecco's Modified Eagle Medium (DMEM) supplemented with phenol red, Glutamax I, and 10% (vol/vol) fetal calf serum (FCS), at 37°C in a 5% $CO_2$ atmosphere. U2OS cells were cultured in McCoy’s 5A medium supplemented with phenol red and 10% (vol/vol) fetal calf serum. For imaging, cells were seeded in μDish IBIDI (Biovalley) coated with poly-L-lysine. Cells were transiently transfected using Genejuice (Merck) or Lipofectamine 2000 (Invitrogen) according to the manufacturer’s protocol for 24 h prior to imaging.

To generate stable cells lines, a kill curve was first determined using G418 (Gibco). Cells were transfected using Lipofectamine 2000 and were exposed to the chosen concentration of G418 after 24 h. The cells were monitored and the media was changed every 24 to 48 h to ensure a constant concentration of G418.

**Zebrafish**

RedFAST-zGem(1-100)-P2A-greenFAST-zCdt1(1-190) mRNA was injected at a final concentration of 100 ng/μL in one-cell stage embryos. Starting from 4 to 8-cell stage, chorions were removed manually and embryos transferred in a glass beaker containing mineral Volvic water. HMBR and HBR-3,5DOM were then added to a final concentration of 5 μM (each), and embryos were incubated 30 to 60 min at 28°C in the dark.

**Fluorescence microscopy**

Confocal micrographs were acquired on a Zeiss LSM 710 Laser Scanning Microscope equipped with a Plan Apochromat 63×/1.4 NA oil DIC M27 immersion objective or a Plan Apochromat 40×/1.4 NA oil DIC immersion objective, heated stage, and XL-LSM 710 S1 incubation chamber for temperature and $CO_2$ control. Images were acquired using ZEN software and processed in Fiji (ImageJ). Photobleaching measurements were acquired using 2.1 kW/cm$^2$, 3.3 kW/cm$^2$, and 13 kW/cm$^2$ at 488 nm and 4.7 kW/cm$^2$ at 541 nm. In all cases the pixel dwell was 1.27 μsec.

To image the split system, rapamycin was added to a final concentration of 500 nM to monitor the association of the FRB-FKBP homodimer. To measure the dissociation of the FKBP-FKBP homodimer, the cells were first pre-incubated with 100 nM AP1510 for ~2 h then rapamycin was added to a final concentration of 1.1 μM for dissociation. To measure multiple PPIs, the cells were treated in the same way as for the FKBP-FKBP homodimer and the association of the homodimer was either followed by imaging every 5 min or was allowed to incubate and the switch between FKBP-FKBP and FKBP-FRB complex was monitored by the addition of rapamycin.

Fluorescence lifetime imaging was performed using a Leica SP8-X-SMD confocal microscope (Mannheim, Germany) with a 63×/1.4 NA oil immersion objective. HMBR and HBR-3,5DOM were excited at 488 nm and 541 nm, respectively, using a ps-pulsed white light laser tuned at 40 MHz. Time-domain FLIM experiments were performed using a time-correlated single-photon counting system operated by an attached PicoHarp 300 module (PicoQuant, Berlin, Germany). Fluorescence emission was detected using two HyDs in photon counting mode at 470-520 nm and 560-595 nm. At least 1000 photon events per pixel were collected and the lifetime analysis was carried out using SymPhoTime software (PicoQuant, Berlin, Germany).

For SOFI imaging, COS7 cells were plated on glass-bottom 35 mm dishes (P35G-1.5-14-C, MatTek) and transfected with pcDNA3-lyn-SkylanS(11) and MAP4-redFAST using FuGene6 (Promega) according to the manufacturer's protocol. The following day, cells were washed with 37°C HBSS and imaged in HBSS supplemented with 5 μM HBR-3,5DOM. The microscope consisted of a Ti2 microscope body carrying a 100× CFI apo TIRF objective (both Nikon) and equipped with a ZT405/488/561/640rpcv2 dichroic and ET525/50m (SkylanS) and ET575lp (redFAST) emission filters (all Chroma). Excitation light was provided by a LBX-488-200-CSB and LBX-405-100-CSB laser operating at 20% and 1%, respectively, for SkylanS and a LCX-561S-100-CSB laser (all lasers Oxxius) operating at 100% for redFAST imaging. The light was fiber-coupled into the microscope body through a manual TIRF module (Nikon) that was aligned in TIRF mode. Images were acquired with a sCMOS camera (Hamamatsu Orca Flash4.0 v2) operating at 50 Hz. We recorded 1000 images in the red channel followed by 1000 images in the green channel. SOFI analysis was performed using the Localizer package (12). For both the green and red images, the first 100 images were discarded and SOFI analysis was performed using the last 900 images, using "Few" pixel combinations.

For zebrafish imaging, embryos were embedded in low-melting agarose (0.8%) extemporaneously mixed with 5 μM final of HMBR and HBR-3,5DOM. Fluorogen solution was then added above the gellified agarose. Imaging was performed with a CSU-W1 Yokogawa spinning disk coupled to a Zeiss AxioObserver Z1 inverted microscope equipped with a sCMOS Hamamatsu camera and a 25× (Zeiss 0.8 Imm WD: 0.19mm) oil objective. DPSS 150 mW 491 nm and 100 mW 561 nm lasers were used with their corresponding 525/50 and 595/50 bandpass excitation filters to respectively acquire greenFAST and redFAST signal. Quantification was performed by measuring the nuclear signal on fluorescent cells over time. The background value was subtracted for each channel and values were normalized to the maximum value of each FAST signal.
